# The lipid Gb3 promotes germinal center B cell responses and anti-viral immunity

**DOI:** 10.1101/2023.09.23.559132

**Authors:** Pankaj Sharma, Xiaolong Zhang, Kevin Ly, Yuxiang Zhang, Yu Hu, Adam Yongxin Ye, Jianqiao Hu, Ji Hyung Kim, Mumeng Lou, Chong Wang, Quinton Celuzza, Yuji Kondo, Keiko Furukawa, David R. Bundle, Koichi Furukawa, Frederick W. Alt, Florian Winau

## Abstract

Influenza viruses escape immunity due to rapid antigenic evolution, which requires vaccination strategies that allow for broadly protective antibody responses. Here, we demonstrate that the lipid globotriaosylceramide (Gb3) expressed on germinal center (GC) B cells is essential for the production of high-affinity antibodies. Mechanistically, Gb3 binds and disengages CD19 from its chaperone CD81 for subsequent translocation to the B cell receptor (BCR) complex to trigger signaling. Abundance of Gb3 amplifies the PI3-kinase/Akt/Foxo1 pathway to drive affinity maturation. Moreover, this lipid regulates MHC-II expression to increase diversity of T follicular helper (Tfh) and GC B cells reactive with subdominant epitopes. In influenza infection, Gb3 promotes broadly reactive antibody responses and cross-protection. Thus, we show that Gb3 determines affinity as well as breadth in B cell immunity and propose this lipid as novel vaccine adjuvant against viral infection.

**One Sentence Summary:** Gb3 abundance on GC B cells selects antibodies with high affinity and broad epitope reactivities, which are cross-protective against heterologous influenza infection.

---

During an infection or immunization, B lymphocytes enter the germinal center (GC) reaction and differentiate into antibody-secreting plasma or memory B cells (*1*). Within GCs, B cells recognize and acquire antigens based on B cell receptor (BCR) affinities and compete for limiting amounts of T cell help, which is necessary for survival, proliferation, and selection of affinity-matured plasma and memory B cells (*2*). However, some pathogens, such as influenza virus, escape the host immune response since their surface antigens display a high frequency of point mutations (antigenic drift) (*3, 4*). The discovery of human broadly neutralizing antibodies (bnAbs), which are able to neutralize distantly related influenza strains, has spurred vaccination strategies aimed at broader humoral responses (*5–7*).

The quality of selected antibodies is determined by GC B cell intrinsic signaling driven by the BCR, co-receptors like CD19, and via interaction with follicular helper T cells (Tfh) (*8–10*). Although there is evidence that the lipid microenvironment affects BCR signaling and Tfh cell functions, overall, the specific roles of lipids in immunoregulation are still underexplored (*10–12*). In this study, we investigated the function of globotriaosylceramide (Gb3), which has been described as GC B cell marker for many years (*13–16*). Gb3 is a glycosphingolipid that is located in membranes of endothelial cells and tubular epithelial cells in the kidney (*17*). However, in the immune system, Gb3 is abundantly expressed on GC B cells. Despite its phenotypic role as marker CD77, the functional impact of Gb3 on B cell responses had not been addressed.

## Gb3 abundance on GC B cells is vital for antibody affinity maturation

To modulate Gb3 levels, we used mice that lack the key enzymes involved in synthesis (α1,4-galactosyltransferase; A4Galt) or degradation (α-galactosidase A; Gla) of Gb3 (fig. S1A). To calibrate this model for Gb3 abundance, we first immunized WT, *A4galt-*KO, and *Gla-*KO mice with 4-hydroxyl-3-nitrophenylacetyl coupled to ovalbumin (NP-OVA) (Fig. 1A) and determined the number of Gb3-positive leukocytes. Of note, Gb3 expression was restricted to GC B cells (fig. S1, B to E). Whereas *A4galt*-KO mice expectedly failed to express Gb3, WT and *Gla*-KO animals showed ~20% and 50% Gb3^pos^ GC B cells (fig. S1, F and G). Thus, *A4galt*-KO, WT, and *Gla*-KO mice represent Gb3-deficient, homeostatic, and Gb3-abundant conditions, respectively, and hence are a suitable model to test the functional impact of various Gb3 levels on GC responses.

**Fig. 1.**
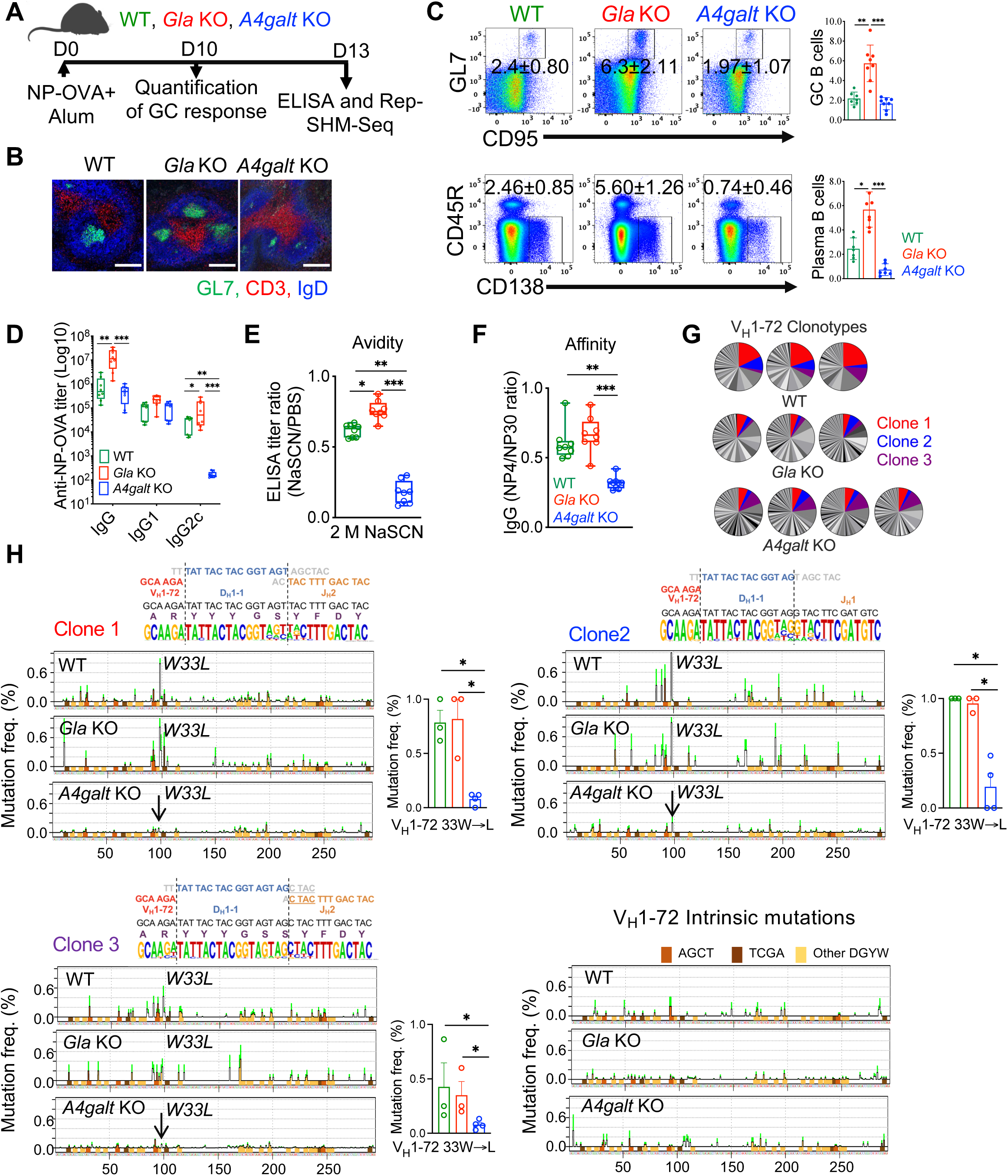
Gb3 promotes the formation of germinal centers and antibody maturation. (A) Diagrammatic representation of the experimental set-up. WT, *Gla*-KO, or *A4galt*-KO mice were immunized with NP-OVA adsorbed on alum. (B) Confocal microscopy of frozen sections from the spleen on day 10 post immunization. IgD (blue: naive B cells), CD3 (red: T cells), and GL7 (green: GC B cells). Scale bar, 200 μm. (C) FACS plots and percentage of GC B cells and plasma cells in the spleen on day 10 post NP-OVA immunization. Each dot represents one mouse (7 or 8 mice per group), and the experiment was repeated three times. (D) Serological analysis of immunoglobulin concentrations of indicated isotypes based on end-point titers using ELISA on day 13 post immunization. Experiment was repeated three times with 8 mice per group. (E) Avidity of NP-OVA-specific IgG was measured by ELISA and expressed as the ratio between the end-point titer values obtained with or without sodium thiocyanate treatment (2 M) on day 13 after immunization. Experiment was repeated twice with 8 mice per group. (F) Antibody affinity assay. Sera were analyzed by ELISA and high-affinity antibodies were measured as a ratio of antibodies binding to BSA conjugated with NP4 (low valency) to NP30 (high valency). Experiment was repeated twice with 8 mice per group. (G) Pie charts representing the distribution of productive V_H_1-72 clonotypes in the GCs of each mouse (3 mice in WT and *Gla*-KO groups and 4 in *A4galt*-KO). Colors indicate the dominant clonotypes found among three different mouse genotypes. (H) SHM profile of V_H_1-72 segments of the dominant clonotypes and intrinsic mutation pattern. Data are plotted as the mutation rate at each nucleotide with representative junctional structure of each IgH clonotype shown at the top, while bar graphs depict individual mice for the frequency of the NP-specific high-affinity mutation (W33L). *p*-values in all graphs were calculated by using Kruskal-Wallis H test with Dunn’s multiple comparison test.

Following NP-OVA immunization, elevated Gb3 abundance in *Gla*-KO mice amplified frequencies of GC B, plasma, and Tfh cells in the spleen and sustained long-lived plasmablasts in the bone marrow (Fig. 1, B and C, and fig. S1, H and I). By contrast, Gb3-deficient *A4galt*-KO animals lacked proper GC organization and exhibited poor differentiation of plasma and Tfh cells (Fig.1, B and C, and fig. S1I). Notably, *Gla*-KO mice showed higher titers of total IgG, whereas Gb3 deficiency abrogated IgG2c class switch (Fig. 1D). To validate the B cell intrinsic function of Gb3, we used mixed bone marrow chimera and found that Gb3-abundant GC B cells outcompeted WT cells and showed an increased proportion of IgG2c-expressing cells (fig. S3, A to C). Additionally, we transferred naïve B cells from our mouse models to muMT-KO animals, which lack mature B cells, and observed that elevated GC B cell numbers and IgG2c class switch were intrinsically Gb3-dependent (fig. S3, D to F).

We next measured antibody maturation by quantifying avidity and affinity of antigen-antibody complexes. Antibodies from *A4galt*-KO mice had low avidity, as indicated by the quick denaturation of antigen-antibody complexes in the presence of NaSCN (Fig. 1E). Furthermore, antibodies from *A4galt*-KO serum showed weak binding to BSA substituted with a low number of NP residues, suggesting impaired affinity maturation (Fig. 1F). To perform an in-depth analysis of the BCR repertoire in GC B cells, we used the recently described Rep-SHM-Seq method to detect specific BCR usage and assess somatic hypermutation (SHM) patterns in enriched clonotypes, where ≥90% nucleic acid sequence similarity in the CDR3 region was used as cutoff to define a clonotype (*18*). To examine heavy chain variable gene segment (V_H_) enrichment, we compared V_H_ usage of GC B cells with naive B lymphocytes. In line with previous studies (*19, 20*), we observed enrichment of NP-specific V_H_1-72 in GC B cells in all immunized mice of the different genotypes (fig. S2). V_H_1-72 rearrangement in GC B cells generated three shared clonotypes among the top five clonotypes in all animal groups (Fig. 1G). The mutation analysis of clonotypes revealed that dominant V_H_1-72 B cell clones from all mice had G-to-T point mutations at residue 98 of CDR1, which corresponds to the mutation W33L that affords high affinity for NP-antigen. While there was no difference in the frequencies of W33L mutations in WT and Gb3-abundant mice, GC B cells lacking Gb3 showed drastically lower frequencies of the high-affinity W33L mutation in all dominant V_H_1-72 clonotypes (Fig. 1H). Altogether, these data demonstrate that Gb3 abundance expands GC B, plasma and Tfh cells, and mice deficient in Gb3 show a lack of antibody affinity maturation.

## Gb3 binds to CD19 for translocation to the BCR complex to trigger signaling

Within germinal centers, centroblasts in the dark zone undergo extensive proliferation as well as somatic hypermutation. Subsequently, centroblasts transition into the light zone to become centrocytes, which undergo affinity-based selection (*8, 21*). Since the transition between dark and light zone is critical for effective humoral responses, we next examined the impact of Gb3 on the composition of centroblasts and centrocytes. Analyzing GC B cells for the centroblast marker CXCR4 and the centrocyte molecule CD83, we found that centroblast/centrocyte transition was severely hampered in the absence of Gb3 (Fig. 2, A and B). This suggests that the lipid Gb3 is crucial for proper positioning of GC B cells in the light zone to allow for productive selection processes to occur.

**Fig. 2.**
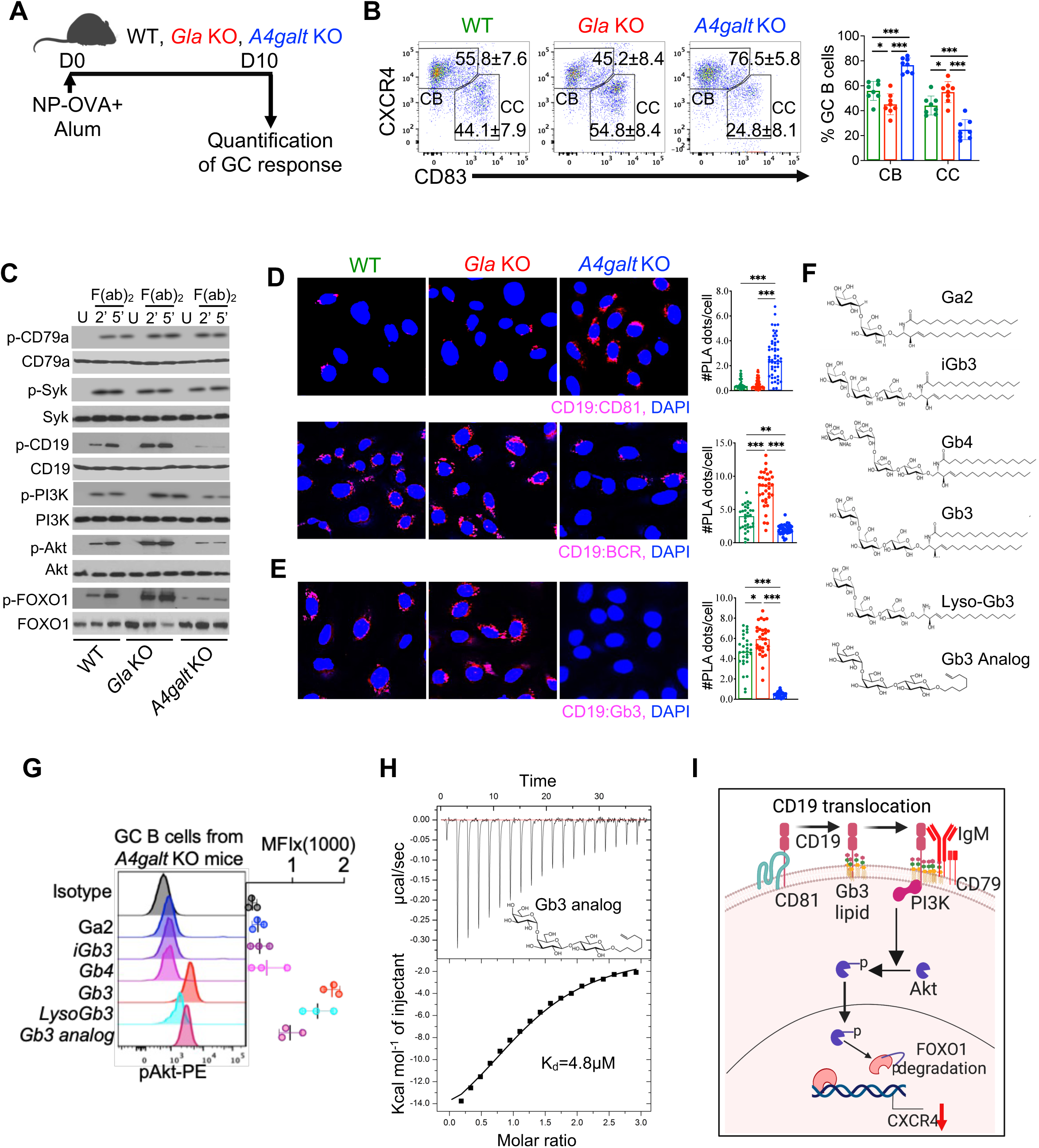
Gb3 binds to CD19 for translocation to the BCR complex and efficient signaling. (A) Diagrammatic representation of the experimental set-up. WT, *Gla*-KO, or *A4galt*-KO mice were immunized with NP-OVA adsorbed on alum. (B) FACS plots and percentages of centroblasts (CB) and centrocytes (CC) in the spleen on day 10 post immunization. Each dot represents one mouse (7 or 8 mice per group), and the experiment was repeated three times. (C) Immunoblot showing BCR and CD19 downstream signaling molecules and transcription factors. Anti-BCR antibodies (F(ab)_2_) were used to stimulate FACS-sorted GC B cells for 2 or 5 minutes. U = unstimulated. (D) Proximity ligation assay (PLA) performed on GC B cells to probe for vicinity between CD81 and CD19 (top panel; blue = DAPI, red = CD19:CD81 PLA signal), and BCR and CD19 (bottom panel; blue = DAPI, red = CD19:BCR PLA signal). In all panels, experiments were repeated at least three times, and PLA signal on more than 30 cells in different fields was calculated for statistical analysis. (E) PLA performed on FACS-sorted GC B cells to probe for proximity between CD19 and Gb3 (blue = DAPI, red = CD19:Gb3 PLA signal). PLA signal was captured by confocal microscopy, and images were processed and analyzed by Image J software. Experiment was repeated at least three times, and PLA signal on more than 30 cells in different fields was calculated for statistical analysis. (F) Structures of different lipids and Gb3 analog used in the study. (G) Phospho-flow to examine the effect of lipid reconstitution on Akt phosphorylation in GC B cells from *A4galt*-KO mice. Histogram overlay and mean fluorescence intensity (MFI) of pAkt staining in GC B cells cultured with different lipids (bottom panel). FACS-sorted GC B cells from *A4galt*-KO mice were seeded with a complex of lipid and BSA for 2h at 37°C (see methods), and Akt phosphorylation was quantified after stimulation of GC B cells with F(ab)_2_. (H) Isothermal titration calorimetry (ITC) to measure the binding between Gb3 and CD19. CD19 and the Gb3 analog were dissolved in PBS, and thermodynamic analysis of their binding was carried out at 25°C on a MicroCal ITC 200 instrument. Top panel: x-axis depicts time, and y-axis represents rate of heat release (μcal/sec). Bottom panel: x-axis represents molar ratio between CD19 and Gb3-analog, and y-axis depicts change in enthalpy. (I) A mechanistic scheme of the effect of Gb3 on the CD19 translocation and BCR downstream signaling pathway. The *p*-values in all graphs were calculated by Kruskal-Wallis H test with Dunn’s multiple comparison test.

GC B cell proliferation in the dark zone in response to BCR activation is mainly regulated by the transcription factor Foxo1 (*22–24*). CD19 positively regulates BCR signaling, and cross-linking of BCR with CD19 reduces its activation threshold (*10, 25*). Furthermore, mice containing loss-of-function mutations at two signaling tyrosine residues of CD19 (Y482F and Y513F) lack selection of antigen-specific antibodies (*26*). Phosphorylation of CD19 activates the PI3K-Akt pathway that leads to Foxo1 phosphorylation and triggers its proteasomal degradation, thus terminating the Foxo1-dependent gene signature that includes CXCR4 expression (*24, 26, 27*). Since we observed accumulation of CXCR4-expressing centroblasts in Gb3 deficiency, we next explored the impact of Gb3 on BCR signaling and downstream activation of the PI3K/Akt/Foxo1 pathway. We isolated GC B cells from our different mouse models and subjected them to BCR ligation with anti-F(ab)_2_ antibodies, prior to analyzing the phosphorylation of signaling molecules using immunoblot. Modulation of Gb3 abundance had no differential effect on phosphorylation in the proximal signal cascade, involving CD79b and Syk (Fig. 2C). However, Gb3-abundant GC B cells showed strong CD19 downstream signaling, markedly higher phosphorylation of Akt and consequently low expression of total Foxo1 (Fig. 2C). In striking contrast, CD19 downstream signaling was compromised in Gb3-deficient GC B cells, as they exhibited diminished phosphorylation of CD19, PI3K, and Akt. The low level of p-Akt in Gb3-deficient GC B cells resulted in impaired phosphorylation and concomitant accumulation of total Foxo1 (Fig. 2C). Taken together, these data indicate that Gb3 activates the CD19-PI3K-Akt axis, leading to Foxo1 degradation, which corresponds to CXCR4 downregulation and efficient centroblast/centrocyte transition.

In resting B cells, CD19 forms a complex with CD81, a tetraspanin required for CD19 trafficking to the B cell surface (*28–30*). Activated GC B cells exhibit pronounced phosphatase activity and transient activation of kinases after BCR cross-linking (*31*), which makes the phosphorylation of signaling molecules a delicate process. This challenge is overcome by the lateral translocation of CD19 from CD81 to the BCR signaling domain in activated B cells (*12, 29, 32*). Since Gb3 abundance drives strong CD19 downstream signaling, we hypothesized that the lipid Gb3 might facilitate CD19 disengagement from CD81 for translocation to the BCR nanocluster to form a functional signaling complex. To this end, we explored the interaction of BCR and CD81 with CD19 using proximity ligation assays (PLA). Gb3-deficient GC B cells displayed strikingly high PLA signal for CD81-CD19 proximity, while in a complementary fashion, BCR-CD19 proximity was markedly reduced compared to WT and *Gla*-KO mice (Fig. 2D). By contrast, BCR-CD19 PLA signal was significantly augmented in *Gla*-KO GC B cells, indicating that Gb3 abundance dissociates CD19 from CD81 and amplifies the spatial relationship between CD19 and the BCR (Fig. 2D). In the BCR downstream signaling cascade, PI3K is physically recruited to the cytoplasmic tail of CD19. Performing PLA, GC B cells abundant in Gb3 showed increased interaction between CD19 and PI3K after BCR cross-linking (fig. S4A). Most notably, the PLA signal in *A4galt*-deficient GC B cells was virtually abolished (fig. S4A), suggesting that the lipid Gb3 is critically required for PI3K recruitment to CD19. Furthermore, PLA between Gb3 and CD19 revealed a direct interaction between the lipid and the co-receptor (Fig. 2E), which indicates that direct interaction with Gb3 promotes the clustering of CD19 with the BCR complex and consequently amplifies CD19 downstream signaling after BCR stimulation.

Next, we tested if the lipid Gb3 is able to exert a direct effect on GC B cell stimulation and whether the carbohydrate headgroup and/or the lipid tail are required for its activation-promoting properties. For this purpose, we cultured Gb3-deficient GC B cells in the presence of galabiaosylceramide (Ga2), Gb3, globotriaosylsphingosine (Lyso-Gb3), isoglobotriaosylceramide (iGb3), globotetraosylceramide (Gb4), or a Gb3-sugar analog for two hours. Ga2, iGb3, and Gb4 share a similar ceramide backbone like Gb3 and only differ in their carbohydrate headgroups. By contrast, Lyso-Gb3 and the Gb3 analog contain the same sugar headgroup as Gb3 (Fig. 2F). Subsequently, we measured phosphorylation of Akt after BCR cross-linking using phospho-flow (Fig. 2G). Exogenous addition of Gb3, Lyso-Gb3, or the Gb3-sugar analog restored Akt activation in *A4galt*-deficient GC B cells, albeit at different levels (Fig. 2G). By contrast, Ga2, iGb3, and Gb4 had no effect on B cell activation. These results demonstrate the immediate impact of Gb3 on B cell stimulation based on its ability to reconstitute Gb3-deficient GC B cells. Furthermore, Gb3-mediated activation of BCR downstream signaling proved to depend on the lipid’s trisaccharide structure and not the ceramide lipid tail.

We then analyzed direct interaction between CD19 and different lipids using a protein lipid overlay (PLO) assay. PVDF membranes were loaded with different concentrations of lipids, and in accordance with our phospho-flow data, Gb3 strongly bound to CD19, followed by Lyso-Gb3 and Gb3 analog (fig. S4B). Of note, we did not see any CD19 binding with Ga2, iGb3, or Gb4. Next, to quantify binding affinity of Gb3 and CD19, we used isothermal titration calorimetry (ITC), in which the lipid is injected into a reaction chamber containing recombinant CD19 in solution. Since the titration of lipid into aqueous phase is challenging in ITC and the CD19-Gb3 interaction depends on the carbohydrate headgroup, we used the Gb3 analog with a short alkyl chain for the ITC assay. Binding data revealed a specific interaction between Gb3 and CD19 with a stoichiometry of 1:1 and a dissociation constant of 4.8 μM (Fig. 2H). Taken together, these results demonstrate that Gb3 binds to CD19 to enable its translocation to the BCR complex for efficient downstream signaling (Fig. 2I).

## Gb3-mediated IgG2c isotype switch is dependent on type I interferon signaling

Since type I interferon is critical for IgG2c class switching (*33, 34*), and as *Gla*-KO mice had elevated titers of IgG2c, we next asked whether Gb3-dependent IgG2c induction is regulated via signaling through interferon-α receptor (IFNAR1). Because type I interferon signaling is important for GCs to form (*35*), we aimed at testing the impact of IFNAR1 specifically on IgG2c class switching following the initial phase of GC development. To this end, we injected anti-IFNAR1 blocking antibodies to mice with ongoing GC reactions at day 6 and 8 post immunization (fig. S5A). Indeed, the amplified GC B cell response in Gb3-abundant animals was not affected by anti-IFNAR1 treatment (fig. S5B). By contrast, IFNAR1 blocking reduced the total IgG titers in *Gla*-deficient mice to WT levels, and IgG2c production was virtually abrogated (fig. S5C). To further validate the impact of IFNAR signaling on Gb3-mediated IgG2c class switch, we used *Gla*-deficient and *Gla*^-/-^/*Ifnar*^-/-^ (double knock-out, DKO) mice, and quantified IgG1- and IgG2c-positive GC B cells after NP-OVA immunization. *Gla*-deficient and DKO mice showed comparable frequencies of IgG1-expressing cells, whereas IgG2c-positive GC B cells were virtually lacking in the absence of functional IFNAR signaling in DKO mice (fig. S5, D and E). We next analyzed the titers of IgG1 and IgG2c antibodies in serum and found that IgG2c induction was specifically dependent on IFNAR signaling as we observed a significantly reduced ratio of serum IgG2c to IgG1 in DKO animals (fig. S5F).

Furthermore, we mechanistically explored the effect of Gb3 on type I interferon signaling and examined a possible molecular interaction between Gb3 and IFNAR1 on GC B cells, using proximity ligation assays (PLA). Analysis based on confocal microscopy revealed elevated PLA signal for Gb3 and IFNAR1 on GC B cells isolated from *Gla*-KO mice compared to WT, while GC B cells from *A4galt*-deficient mice lacking Gb3 failed to produce any PLA signal (fig. S5G). In order to functionally investigate Gb3-mediated IFNAR1 signaling, we stimulated FACS-sorted GC B cells with IFNα and assessed the activation of downstream kinases. Immunoblot analysis showed drastically augmented phosphorylation of Tyk2, JAK1, STAT1, and STAT2 in GC B cells from Gb3-abundant mice when compared to WT (fig. S5H). In sharp contrast, GC B cells from Gb3-deficient animals displayed strikingly diminished activation of the respective kinases (fig. S5H). Taken together, these data demonstrate that Gb3 interaction with IFNAR1 modulates the type I interferon signaling cascade and facilitates the selection of the IgG2c class of antibodies.

## Gb3 drives B cell repertoire diversity

A central key to effective immunity is the generation of broad antibody responses, which are not limited to the most prominent antigenic determinants, but also recognize subdominant epitopes. Interclonal competition and repetitive mutation and selection processes increase affinity and impair clonal diversity (*36*). Since Gb3-abundant mice exhibited higher numbers of centrocytes, a cell population that undergoes selection in the light zone of GCs, we then asked whether high levels of Gb3 help with the selection of B cell clones reactive against diverse epitopes. For this purpose, we first performed rarefaction analysis of our Rep-SHM-Seq libraries to estimate for species richness, where each clonotype is defined as a unique species (Fig. 3, A and B). Our data show that, while there was no significant difference between WT and *A4galt*-KO, B cell clones from *Gla*-deficient animals showed higher diversity (Fig. 3B).

**Fig. 3.**
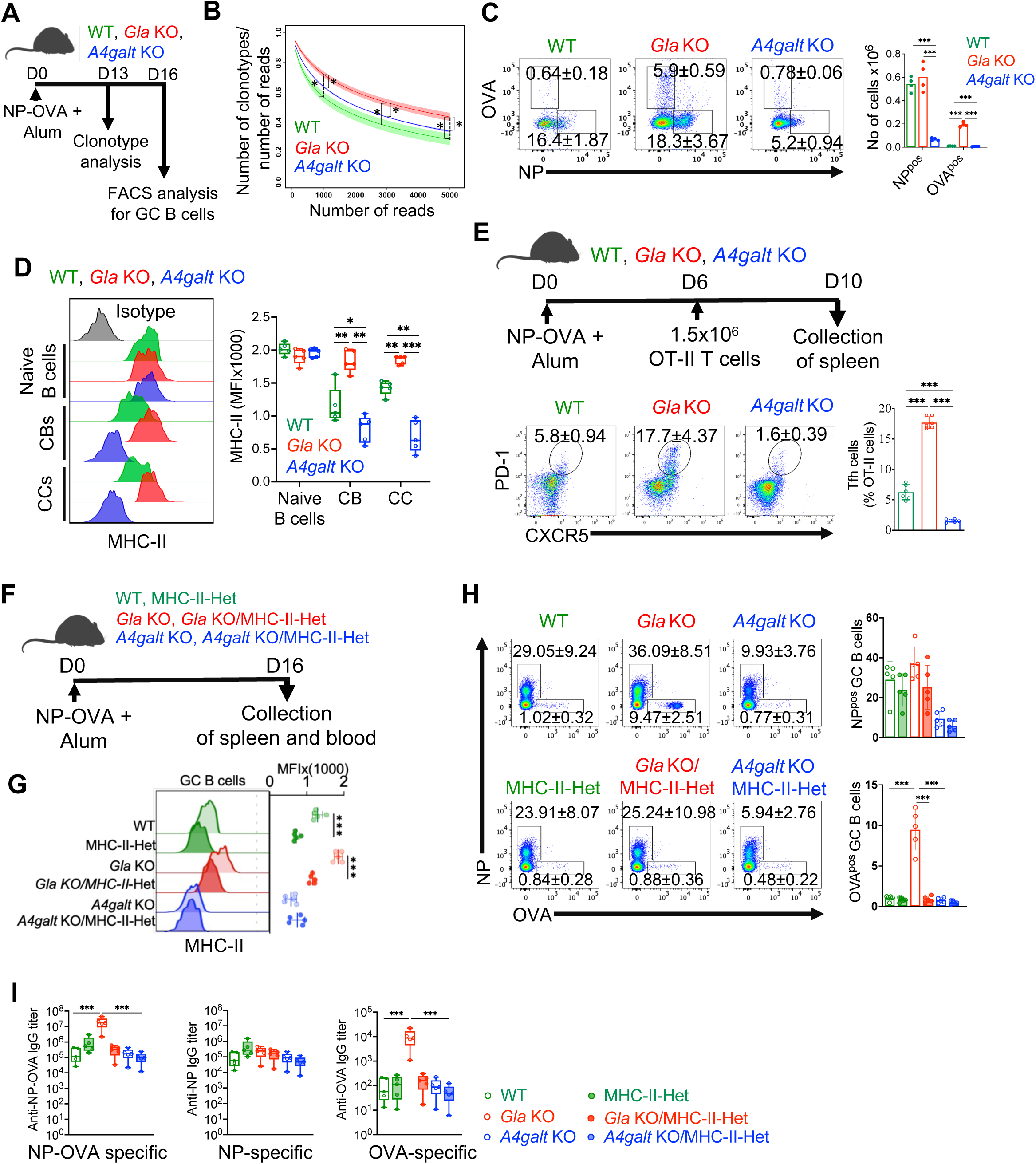
Gb3 facilitates the selection of subdominant epitopes. (A) Schematic representation of the experimental set-up. WT, *Gla*-KO, or *A4galt*-KO mice were immunized with NP-OVA adsorbed on alum. (B) Rarefaction curves showing the clonotype diversity for the three genotypes (green: WT, red: *Gla*-KO, blue: *A4galt*-KO). The solid dark curves and light shades show the mean and SEM (standard error of the mean) of the calculated diversity for each genotype (N=3 for WT and *Gla*-KO, and N=4 for *A4galt*-KO). Student’s t-tests were used to test the difference in diversity across different genotypes at the number of reads 1000, 3000, and 5000 (n=3 for WT and *Gla*-KO, n=4 for *A4galt*-KO). (C) FACS plots and percentages of OVA-FITC^+^ and NP-PE^+^ GC B cells in the spleen on day 16 post immunization. Each dot represents one mouse (4 mice per group), and the experiment was repeated at least three times. (D) Histogram overlays depicting the MHC-II expression on naive B cells, centroblasts, and centrocytes as quantified by flow cytometry. Each dot represents a single mouse with 5 mice per group. Experiment was repeated at least three times. (E) Experimental set-up of NP-OVA immunization and OT-II T cell transfer (top panel). FACS plots and percentages of OT-II Tfh cells in the spleen on day 10 post immunization as quantified by flow cytometry (bottom panel). Dots in bar graphs represent individual mice (5-6 mice per group) and the experiment was at least repeated two times. (F) Experimental set-up of NP-OVA immunization using different mouse models. (G) MHC-II expression on GC B cells on day 16 after immunization. Data are plotted as mean fluorescence intensity (MFI) as quantified by flow cytometry (n=5 mice per group). (H) FACS plots and percentages of OVA-FITC^+^ and NP-PE^+^ GC B cells in the spleen on day 16 post immunization. The experiment was repeated three times (5 mice per group). (I) Serological analysis of indicated antigen-specific immunoglobulin concentrations as detected by ELISA on day 16 post immunization. The experiment was repeated three times (5 mice per group). The *p*-values in graphs C to K were calculated by Kruskal-Wallis H test with Dunn’s multiple comparison test.

Next, we investigated whether higher clonal diversity in Gb3-abundant mice represents the selection of GC B cells reactive with subdominant epitopes. Using the NP-OVA system for immunization, NP serves as the dominant antigenic determinant and OVA as subdominant epitope. Detecting GC B cell reactivities using fluorochrome-coupled NP or OVA, analysis by flow cytometry showed that *Gla*-KO mice displayed a sizable population of OVA-reactive B cells, while WT and Gb3-deficient animals only selected for NP-binding BCRs (Fig. 3C). Moreover, compared to WT and *A4galt*-KO, Gb3-abundant mice showed strikingly increased IgG titers specific for OVA (fig. S6, A and B). Further, we were able to detect OVA-specific plasma cells in the bone marrow of *Gla*-deficient mice at day 60 post immunization, highlighting the lasting humoral response to subdominant epitopes that is mediated by Gb3 abundance (fig. S6C). To test whether the effect of Gb3 on the selection of subdominant epitopes is B cell-intrinsic, we used our B cell transfer set-up to muMT-KO mice that lack mature B lymphocytes (fig. S6D). In comparison to the other experimental groups, we observed a higher frequency of OVA^pos^ GC B cells as well as elevated anti-OVA IgG titers in mice that received *Gla*-KO B cells (fig. S6, E and F), corroborating the B cell-intrinsic function of Gb3-mediated selection of B cell reactivity toward subdominant epitopes.

The expansion of Tfh cells and selection of diverse B cell clones depends on antigen presentation by GC B lymphocytes (*37*). Thus, taking into account that GC B cells show a wide expression range of MHC-II (*38*), we next investigated whether Gb3 affects MHC-II levels as well as antigen presentation to Tfh cells. In line with previous studies (*38*), we found lower MHC-II expression on WT centroblasts than on centrocytes and naive B cells. However, mice rich in Gb3 maintained high amounts of MHC-II on centroblasts as well as centrocytes (Fig. 3D). In sharp contrast, Gb3-deficient GC B cell subsets displayed a strikingly lower surface level of MHC-II (Fig. 3D). We then injected CD45.1-congenic OVA-specific CD4 T cells (OT-II) to NP-OVA immunized mice and quantified the generation of Tfh cells (Fig. 3E). Compared to WT, the striking increase of OVA-specific Tfh cells in Gb3-abundant mice was virtually abrogated in the absence of Gb3 (Fig. 3E). The density of MHC-II/peptide complexes on antigen-presenting GC B cells during their interaction with Tfh cells has been described to regulate B cell selection or entry into the GC reaction (*39–41*). Since we observed an extraordinary number of OVA-reactive GC B cells following hapten-based immunization in Gb3-abundant mice, we then addressed whether the selection of GC B cells reactive with subdominant epitopes is dependent on Gb3-regulated MHC-II expression levels. To this end, we compared the GC response in our mouse models expressing MHC-II from a homozygous or haplosufficient gene locus (Fig. 3F). Consistent with prior work (*41*), MHC-II haplosufficiency halved MHC-II surface expression on GC B cells in WT and Gb3-abundant mice, whereas the effect was negligible in Gb3 deficiency which is characterized by overall low amounts of MHC-II (Fig. 3G). Following NP-OVA immunization, the amplified frequencies of GC B and Tfh cells in *Gla*-deficient animals were reduced upon lowered MHC-II expression in heterozygous MHC-II/*Gla*-KO mice (fig. S6, G to I). Most notably, the drastically elevated response to subdominant OVA antigen in Gb3-abundant mice was virtually abolished when MHC-II expression was cut in half (Fig. 3H). These findings were well reflected in serum antibody titers, with anti-OVA IgG in *Gla*-KO/MHC-II-Het animals brought to WT levels, whereas MHC-II haplosufficiency did not affect the antibody response to the dominant hapten NP (Fig. 3I). Thus, the GC response to subdominant epitopes was strictly dependent on Gb3-modulated MHC-II expression. Taken together, these results demonstrate that Gb3 regulates MHC-II-restricted antigen presentation to select Tfh and GC B cells of broad diversity.

## High Gb3 level cross-protects against influenza infection

In viral infection and vaccination, the generation of broadly neutralizing antibodies reactive with subdominant antigens is vital to ensure continued protection in the face of emerging mutant viruses. Thus, we next explored whether Gb3 abundance in GC B cells allows for the selection of diverse antibodies during respiratory flu infection. For this purpose, we infected our mouse models with influenza A virus (H1N1, strain PR8) and monitored the cellular and humoral immune responses. In contrast to *A4galt*-deficient mice, intranasal infection with 10^2^ plaque forming units (PFU) of H1N1 triggered a strong GC response in mediastinal lymph nodes in *Gla*-KO animals (fig. S7, A and B), followed by increased titers of total IgG and IgG2c antibodies when Gb3 was abundant (fig. S7C). The viral antigen hemagglutinin (HA) consists of a globular head, which largely contains the dominant antigenic sites, and a head-supporting stalk domain that harbors subdominant epitopes. Interestingly, Gb3-abundant mice exhibited strikingly increased titers of serum antibodies that reacted with the HA stalk region (fig. S7D), suggesting an amplified response to subdominant antigenic determinants mediated by Gb3. Moreover, *Gla*-deficient animals showed increased antibody titers recognizing recombinant H3, the most distantly related HA molecule compared to H1 (fig. S7E). Next, we used live virus to examine the neutralization capacity of serum antibodies derived from our different mouse models. Compared to WT and *A4galt*-KO animals, neutralization of H1N1 in vitro was augmented using serum from Gb3-abundant mice (fig. S7F). Notably, antibodies from *Gla*-deficient animals also allowed for neutralization of a distantly related heterologous influenza virus (H3N2, strain HK68 X-31), demonstrating Gb3-mediated induction of cross-reactive antibodies (fig. S7G).

Lastly, we performed in vivo challenge experiments, using H1N1 for priming followed by secondary infection with either H1N1 or H3N2 (fig. S7H). Compared to WT, *Gla*-KO mice showed protection against both influenza strains, as reflected by reduced weight loss and low viral burden in the lung (fig. S7, I to L). By contrast, *A4galt*-deficient mice suffered from significant weight loss and failed to control pulmonary viral load (fig. S7, I to L). Interestingly, only mice with high Gb3 levels were able to virtually eradicate H3N2 viral burden in the lungs (fig. S7L), demonstrating Gb3-mediated cross-protection. Altogether, these data indicate that Gb3 in GC B cells drives the recognition of the HA stalk region, development of cross-reactive antibodies, and heterologous protection against influenza infection.

## Exogenous Gb3 is a potent adjuvant for influenza vaccination

Based on our results describing the immunological functions of Gb3 in GC B cells, we next aimed at developing exogenous Gb3 as potential adjuvant for B cell responses upon vaccination. First, we investigated whether administration of synthetic Gb3 is able to reach GC B cells. For this purpose, we immunized *A4galt*-KO mice with emulsified Gb3 in combination with NP-OVA (fig. S8A). Ten days after immunization, we detected ~20% Gb3^pos^ GC B cells in the draining lymph node (fig. S8B). In addition, Gb3-treated mice showed increased total IgG, IgG2c, as well as Akt-phosphorylation in activated GC B cells (fig. S8, C and D). These results demonstrate that injected Gb3 is physically acquired by GC B cells and drives their activation.

Thus, to test the adjuvant effect of Gb3 in anti-viral vaccination, we immunized WT mice with recombinant hemagglutinin (rHA) alone, or in combination with either Gb3 or the well-established adjuvant alum (Fig. 4A). In a prime-boost model, both Gb3- and alum-formulated rHA induced robust antibody responses (Fig. 4, B to D). However, mice immunized with Gb3-formulated rHA showed the strongest production of the IgG2c class of antibodies (Fig. 4D), higher avidity of antigen-antibody complexes (Fig. 4E), and dramatically elevated titers of antibodies reactive with the HA stalk (Fig. 4F). While both alum- and Gb3-adjuvanted animals had robust memory B cell responses, plasma cells secreting IgG2c were exclusively present in the Gb3-adjuvanted group at day 60 post immunization (Fig. 4G). Moreover, the antibodies isolated from the Gb3-adjuvant group bound more effectively to rHA molecules from group 1 (H1 and H5) as well as group 2 (H3 and H7) of influenza viruses (Fig. 4H). Whereas Gb3- and alum-adjuvanted groups had similar H1N1 (PR8) neutralization titers, only Gb3 treatment facilitated efficient neutralization of the group 2 virus H3N2 (X-31) (Fig. 4, I and J). These data demonstrate that exogenous Gb3 acts as a powerful adjuvant and promotes the production of cross-reactive antibodies.

**Fig. 4.**
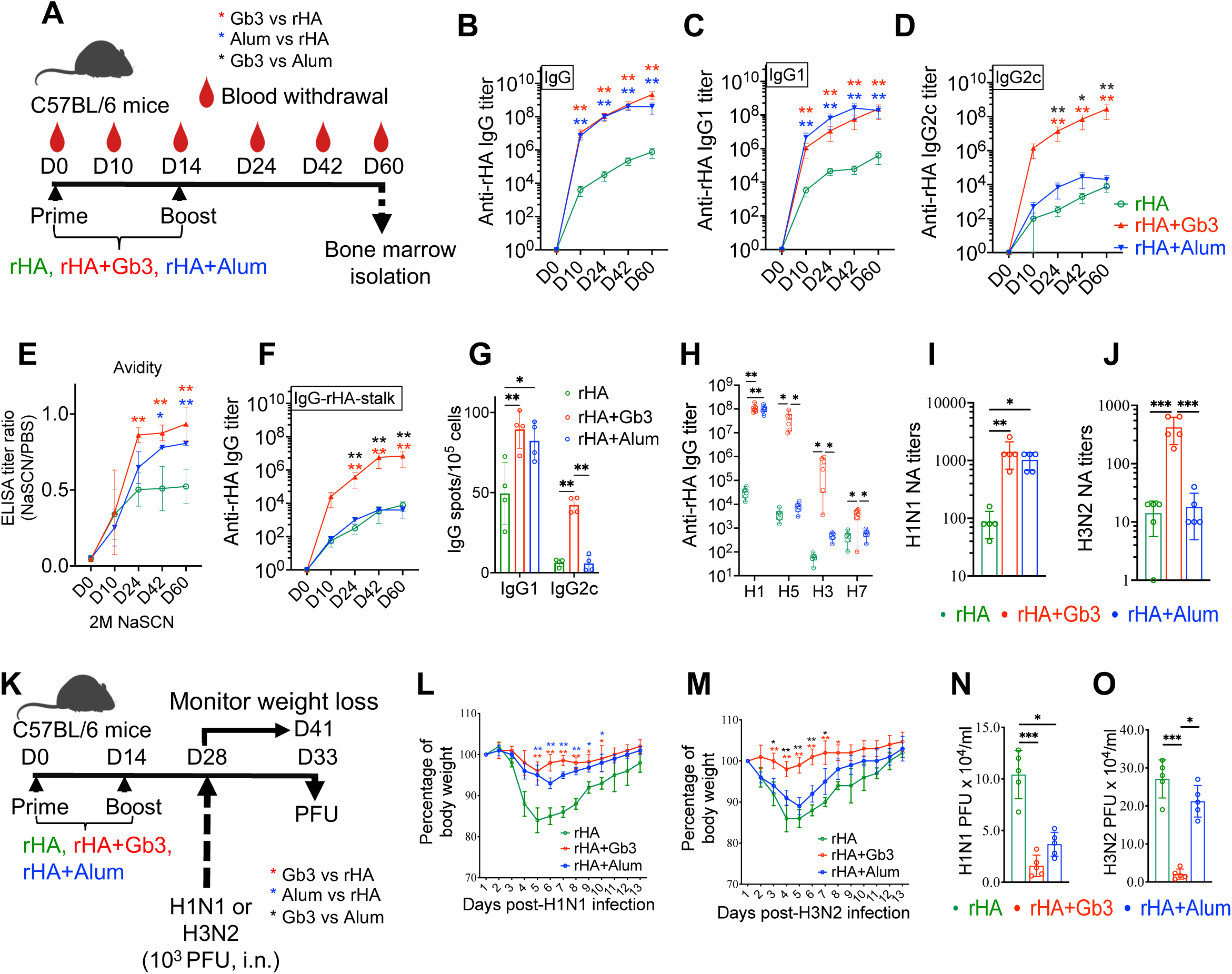
Exogenous Gb3 broadens the antibody response in influenza infection. (A) Experimental set-up depicting animal groups, timeline of immunization, and sample collection schedule for rHA immunization. (B to D) Serological analysis of immunoglobulin concentrations of indicated isotypes specific for rHA was performed by ELISA. Graphs represent mean±SD with 5 mice in each group. The experiment was repeated three times. (E) Avidity of rHA-specific IgG was measured by ELISA and expressed as the ratio between the end-point titer values obtained with or without sodium thiocyanate treatment (2 M). Graphs represent mean±SD with 5 mice in each group. (F) Serum concentrations of anti-rHA stalk immunoglobulins were detected by ELISA. Graphs represent mean±SD with 5 mice in each group. (G) Enzyme-linked immunosorbent spot (ELISPOT) showing IgG1- and IgG2c-producing plasma cells recognizing rHA on day 60 post immunization. Dots in bar graphs represent individual mice (n=4 for each group), and the experiment was repeated three times. (H) Serum concentrations of immunoglobulins against rHA of heterotypic influenza strains were detected by ELISA. Dots in bar graphs represent individual mice (n=6 for each group), and the experiment was repeated three times. (I and J) H1N1 or H3N2 neutralizing antibody (NA) titers in serum on day 28 post rHA immunization. Serum concentrations of immunoglobulins against rHA of heterotypic influenza strains were detected by ELISA. Dots in bar graphs represent individual mice (n=5 for each group), and the experiment was repeated at least three times. (K) Experimental set-up depicting immunization and infection regimen. Immunized mice were challenged intranasally with 10^3^ PFU of either H1N1 or H3N2 on day 28. PFU = plaque forming units; i.n. = intranasal. (L and M) Weight loss curve following H1N1 (PR8) or H3N2 (HK68, X-31) infection. Data shown in graphs represent three independent experiments with 5 mice in each group, and the experiment was repeated twice. (N and O) H1N1 or H3N2 PFUs in lungs on day 5 post infection. Dots in bar graphs represent individual mice, and the experiment was repeated twice. The *p*-values were calculated by Kruskal-Wallis H test with Dunn’s multiple comparison test, and in graph l and m, *p*-values of ≤0.05 and ≤0.01 were marked with * and **, respectively.

Next, we investigated if Gb3-immunized mice were broadly protected against influenza infections with distantly related viral strains. To this end, we infected vaccinated animals with either H1N1 or H3N2 and monitored weight loss and virus multiplication in the lung (Fig. 4K). Only Gb3-adjuvanted rHA provided protection from both viral strains as characterized by reduced weight loss (Fig. 4, L and M) and improved clearance of viruses in the lung (Fig. 4, N and O).

Ultimately, we aimed at translating our findings on Gb3 as an adjuvant to the human system. First, we analyzed whether Gb3 is able to interact with CD19 to drive BCR downstream signaling in human GC B cells. For this assay, we sorted CD77^pos^ and CD77^neg^ GC B cells from human tonsils and in parallel reconstituted CD77^neg^ cells with exogenous Gb3 (fig. S9A). Performing PLA, we found a strong signal in confocal microscopy, indicative of proximity between Gb3 and CD19 in human GC B cells (fig. S9B). Functionally, after BCR cross-linking, CD77^pos^ B cells displayed higher phosphorylation of Akt and degradation of Foxo1 than CD77^neg^ cells. Furthermore, treatment with Gb3 was able to reconstitute effective BCR downstream signaling in CD77^neg^ GC B cells (fig. S9C).

We next compared the adjuvant effect of Gb3 with the clinically used adjuvant alum, using human tonsil organoids as previously described (*42*). Dissociated tonsil cells were cultured in vitro to form organoids and then stimulated with antigens and adjuvants (fig. S9D). Compared to stimulation with rHA alone, Gb3 improved GC responses as well as plasma cell differentiation (fig. S9E). In addition, Gb3 triggered a total anti-rHA IgG1 titer higher than rHA and alum, reminiscent of our findings in the murine model regarding IgG2c production (fig. S9F). Taken together, these results demonstrate that the lipid Gb3 can activate CD19 downstream signaling in human B cells and has great potential as an adjuvant in the human system, involving IgG1-based adjuvanticity.

## Discussion

Beyond its use as a mere marker molecule, we introduce herein novel functions of the lipid Gb3 that regulate the GC response at multiple levels (fig. S9G).

i. *CD19 translocation to the BCR complex, amplified downstream signaling, and centroblast/centrocyte transition.* CD19 is a central signaling hub in B cell activation and its physical recruitment of PI3K is critical for signaling along the Akt/Foxo1 pathway (*43*). Foxo1 in turn is a major transcription factor that dominates the gene expression maintaining the dark zone of the GC, and consequently, PI3K activation and Foxo1 degradation are a prerequisite for centroblast transition to the light zone (*22, 24*). In this context, our data show that Gb3 promotes BCR downstream signaling and amplifies activation of the PI3K/Akt/Foxo1 pathway. An increase of Gb3-mediated Foxo1 degradation allows for efficient centroblast/centrocyte transition, thus driving the overall dynamic of the GC reaction and the opportunity for subsequent selection processes in the light zone. Activation of CD19 requires its dissociation from the resting CD81 complex and subsequent translocation to an active BCR nanocluster (*12, 29*). However, mechanisms controlling lateral translocation of CD19 in activated B cells are not clearly understood. Our work revealed that the glycolipid Gb3 binds to CD19, and this idiosyncratic lipid-protein interaction drives lateral translocation of CD19 to the BCR complex to facilitate the signaling cascade.
ii. *MHC-II presentation, Tfh cell differentiation, and subdominant epitope selection.* B cells equipped with a high-affinity BCR are most successful in taking up antigen presented by FDCs in the light zone, which makes them potent antigen-presenting cells for activation of Tfh cells recognizing MHC-II (*39*). Tfh cells in turn provide critical signals for GC B cell selection (*8, 9*). Thus, the variety of the Tfh cell repertoire is mirrored in the selection of diverse GC B cell clones, including the ones reactive with subdominant epitopes (*37*). In this context, we found Gb3-dependent regulation of MHC-II on GC B cells and that high Gb3 abundance promoted Tfh cell differentiation. Using the NP-OVA system, proliferation of OVA-reactive Tfh cells corresponded to the expansion of GC B cells recognizing OVA as subdominant epitope when Gb3 was abundant. Furthermore, MHC-II haplosufficient mice lacked any subdominant epitope reactivity, which highlights an important role of Gb3 in MHC-II-restricted antigen presentation and the coordinated interaction between GC B and Tfh cells.

Infecting either *Gla*-deficient mice or vaccinated animals with influenza virus, we discovered three Gb3-mediated hallmarks in flu infection: IgG2c production, recognition of the HA stalk, and generation of cross-reactive antibodies that protect against diverse viral strains. IgG2c is important since it proved to be the most effective isotype for influenza protection (*44*). This is based on the binding capacity of IgG2c to Fc receptors on myeloid cells, which amplifies antibody-dependent cell-mediated cytotoxicity and clearance of infected host cells (*45, 46*). In contrast to the influenza virus HA head, which represents the main target of antibody responses, the stalk region harbors subdominant epitopes, which are less frequently mutated and therefore able to sustain effective immunity (*7, 47*). Thus, novel vaccine strategies aim at generating broadly neutralizing antibodies that are independent of seasonal antigenic drift and protect against a large range of viral strains (*48*). Interestingly, using exogenous Gb3 for vaccination, we achieved cross-reactive, heterologous protection against influenza viruses, suggesting that Gb3 could serve as a future adjuvant toward a universal vaccine that might ultimately eradicate the flu.

## Author contributions

P.S. and F.W. designed the study, performed data analysis, and wrote the manuscript. P.S., X.Z., K.L., Y.Z., Y.H., J.H.K., M.L., and C.W. performed experiments. Y.Z., A.Y.Y., Q.C., J.H., and F.W.A. analyzed BCR sequencing data. D.R.B. synthesized the Gb3 analog. Y.K., K.F., and K.F. generated *A4galt*-KO mice. All authors reviewed the manuscript.

## Acknowledgements

We are grateful to K. Arnett and the Center for Macromolecular Interactions at Harvard Medical School for assistance with ITC experiments. This work was supported by National Institutes of Health grants R01 AI136939 to F.W.

## Data and material availability

All data are available in the manuscript or the supplementary material.

## Supplementary material

Materials and Methods

Figs. S1 to S9

Table S1

References and notes (49-53)

## Materials and Methods

### Reagents

RPMI medium 1640, ethylenediaminetetraacetic acid (EDTA), β-mercaptoethanol, penicillin and streptomycin, sodium pyruvate, L-glutamine, and TMB Substrate Solution were purchased from Thermo Fisher Scientific (Waltham, MA, USA). Fetal bovine serum (FBS), acetylated trypsin, bovine serum albumin (BSA), sodium thiocyanate (NaSCN), phosphate-buffered saline (PBS), PLA probe maker kit and Duolink® in situ detection reagents were procured from Millipore Sigma (Burlington, MA, USA). RPMI medium 1640 was supplemented with 10% FBS, penicillin (100 U/ml) and streptomycin (100 μg/ml), sodium pyruvate (1 mM), L-glutamine (2 mM), β-mercaptoethanol (50 µM), and HEPES (100 mM). Gb3 was purchased from Matreya LLC (State College, PA, USA), while other lipids used in the study were purchased from Avanti Polar Lipids (Alabaster, AL, USA). NP-OVA and NP-PE were procured from SouthernBiotech (Birmingham, AL, USA). B cell enrichment kit was purchased from Miltenyi Biotec (Bergisch Gladbach, Germany). Recombinant mouse CD19, hemagglutinin proteins from H1N1 (PR8), H3N2 (HK68, X-31), H5N1 (A/Vietnam/1194/2004) and H7N7 (A/chicken/Netherlands/1/03), and hemagglutinin stalk protein (HA2 subunit) were purchased from Sino Biologicals Inc (Beijing, China). Influenza viruses (H1N1; PR8 and H3N2; HK8) and Madin Darby Canine Kidney (MDCK) cell line were procured from ATCC. Alkaline Phosphatase Substrate Kit III for ELISPOT was procured from Vector Laboratories (Newark, CA, United States). Nitrocellulose membranes, PVDF membranes, 12% precast polyacrylamide gel, X-ray films, and Laemmli buffer were purchased from BioRad laboratories (Hercules, CA, USA). Chemiluminescence substrate (ECL plus) for immunoblots and BCA protein assay kit were procured from Thermo Fisher Scientific (Waltham, MA, USA), and receptor destroying enzyme (RDE II) for hemagglutination inhibition (HI) test was purchased from Denka Seiken (Tokyo, Japan).

### Antibodies

Anti-mouse antibodies against CD3 (145-2C11), CD4 (GK1.5), CD45.1 (A20), CD45.2 (104), CXCR5 (L138D7), PD1 (29F.1A12), GL-7 (GL7), CD45R (RA3-6B2), IgG (Poly4053), IgG2c (MRG2c-67), CD103 (2E7), CD11b (M1/70), F4/80 (QA17A29), Gr-1 (RB6-8C5) as well as anti-human antibodies against CD3 (UCHT1), IgD (IA6-2), CD27 (M-T271), CD38 (HB-7), CD77 (5B5), and purified anti-mouse IgM (RMM-1) antibodies were purchased from BioLegend (San Diego, CA, USA). Purified anti-mouse CD19 (MB19-1), PI3K (H.843.0), CD79a (SP18) and pCD79a (polyclonal), anti-human CD19 (HIB19) and IgM (SA-DA4) antibodies were purchased from Thermo Fisher Scientific (Waltham, MA, USA). The following antibodies for western blot analysis were purchased from Cell Signaling Technology (Danvers, MA, USA): anti-mouse CD19 (D4V4B), Akt (40D4), Syk (D3Z1E), PI3K (C73F8), FOXO1 (C29H4), β-actin (13E5), p-Akt (D9E), p-Syk (65E4), p-PI3K (E3U1H), p-FOXO1 (E1F7T), STAT-2 (D9J7L), Jak1 (D1T6W), Tyk2 (E9H4T), p-STAT-1 (polyclonal), p-Jak1 (D7N4Z), p-Tyk2 (D7T8A), and anti-human β-actin (8H10D10). Anti-mouse p-STAT-2 (polyclonal), STAT-1 (STAT1-79), and p-CD19 (polyclonal) were purchased from Thermo Fisher Scientific (Waltham, MA, USA), while anti-mouse CD77 (BGR23) was purchased from Creative Biolabs. Anti-mouse IgG-HRP (polyclonal), IgG1-HRP (polyclonal), IgG2c-HRP (polyclonal) and anti-human IgG1-HRP were purchased from SouthernBiotech (Birmingham, AL, USA). Cytofix/Cytoperm buffer kit and Fc-blocking reagent were purchased from BD Biosciences (Franklin Lakes, NJ, USA).

### Mice

C57BL/6, B6.SJL (B6.SJL-Ptprca Pepcb/Boy), muMT-KO (B6.129S2-Ighmtm1Cgn/J), MHC-II-KO (B6.129S2-H2dlAb1-Ea/J), OT-II transgenic mice (B6.Cg-Tg(TcraTcrb)425Cbn/J), *Ifnar1*-KO (B6.129S2-Ifnar1tm1Agt/Mmjax), and *Gla*-KO (B6;129-Glatm1Kul/J) mice were acquired from The Jackson Laboratory. *A4galt*-deficient mice were generated by Koichi Furukawa (*49*). Mice were bred and maintained in the animal barrier facility of Harvard Medical School (HMS). All animal procedures were approved by the IACUC at HMS (approval number IS00001618).

### Immunizations and influenza infection

Age- and sex-matched mice were immunized with 25 µg NP-OVA, or 10 µg of rHA (from H1N1; PR8) either alone or in combination with 50 µg alum in 200 µl PBS. To test the adjuvant effect of Gb3, 10 µl of a 5 mg/ml Gb3 stock (in DMSO) was suspended in saline solution containing antigens and emulsified with 10% Tween 80 for 30 minutes. Thereafter, mice were immunized with lipid-formulated antigens in 200 µl PBS. rHA-immunized mice were challenged with 1 x 10^3^ PFU of either H1N1 (PR8) or H3N2 (HK68, X-31) viruses. In separate infection experiments, WT, *Gla*-KO, and *A4Galt*-KO mice were intranasally primed with 1 x 10^2^ PFU of H1N1, and 24 days post priming were challenged with 1 x 10^3^ PFU of either H1N1 or H3N2 viruses. Mice were monitored for weight loss, and 25% weight loss was considered as endpoint.

### Generation of bone marrow chimeric mice

Mixed bone marrow (BM) chimeric mice were generated following an adaptation of previously described methods (*41, 50*). Briefly, donor mice were sacrificed, and their bones were surgically cleaned from surrounding tissue. Extremities of tibiae and femora were trimmed with sterile scissors, and bone marrow was flushed through a 70 µm nylon mesh strainer (Corning Life Sciences, NY, USA). Cell number and viability was determined by trypan blue exclusion. To generate BM chimera, WT.SJL (F1 generation of CD45.2 C57BL/6 and CD45.1 SJL) were lethally irradiated with 1,100 rad and intravenously received 5 × 10^6^ WT CD45.1 and 2.5 × 10^6^ CD45.2 BM cells from either C57BL/6, *Gla*-KO, or *A4galt*-KO mice (ratio of CD45.1:CD45.2 = 2:1). Mice were given drinking water containing 1% neomycin for one week and were used 6 weeks after BM transplantation.

### Adoptive transfer of B cells to muMT-KO mice

6-8-week-old WT, *Gla*-KO, and *A4galt*-KO mice were sacrificed, and total splenic B cells were purified using a negative B cell isolation kit (Miltenyi Biotec). Purity of enriched B cells was determined by flow cytometry, and recipient muMT-KO mice were injected with 1 × 10^7^ B cells intravenously. B cell-reconstituted muMT-KO mice were immunized 7 days after transfer with NP-OVA adsorbed on alum as described above, and GC B cell and antibody responses were quantified on day 21 after B cell transfer.

### ELISA, measurement of antibody affinity and avidity

Total IgG, IgG1, and IgG2c antibodies specific for NP-OVA, OVA, or rHA were quantified using ELISA. High-binding flat-bottom 96-well plates (Corning Life Sciences; NY, USA) were coated with 1 µg/ml NP-OVA, OVA, or rHA in PBS, incubated overnight at 4°C and blocked with PBS + 2% BSA for 1h at room temperature (RT). Subsequently, sera from vaccinated or H1N1-primed mice were added with an initial dilution of 1:100 and 1:4 serial dilutions in PBS + 1% BSA, prior to incubation for 2h at RT. Plates were then washed and incubated for 1h at RT with HRP-conjugated anti-mouse IgG, IgG1, and IgG2c antibodies. Following incubation, plates were washed again and developed with tetramethylbenzidine for 5 min, then stopped with 2 N H_2_SO_4_. The optical density was read at 450 nm using a Versamax microplate reader with SoftMax Pro Version 5 (both from Molecular Devices; San Jose, CA, USA), and endpoint titers were calculated with a cutoff three times the optical density over background. To test the IgG1 released by human tonsil cells, rHA-coated plates were incubated with culture supernatant with an initial dilution of 1:25 and 1:4 serial dilutions. For measuring antibody affinity, plates were coated with either NP30-BSA or NP4-BSA (10 μg/ml). The serum titers of IgG reactive with each conjugate were determined as above. Results were presented as ratios of IgG titers against NP4-BSA versus NP30-BSA. For assessing antibody avidity, plates were incubated 15 min with 2 M sodium isothiocyanate before the addition of HRP-conjugated anti-mouse IgG antibodies. Avidity was expressed as the LogEC50 ratio of corresponding plates treated with or without sodium isothiocyanate.

### ELISPOT

To quantify the antigen-specific antibody-secreting plasma cells by ELISpot assay, 96-well ELISpot plates (Millipore, Burlington, MA, USA) were coated for 1h at room temperature (RT) with 50 μl of 8 μg/ml goat-anti-mouse IgG (KPL, Gaithersburg, MD, USA) for assessing total IgG, or 50 μl containing 20 μg/ml purified rHA proteins from H1N1 (PR8) or H3N2 (HK68, X-31) influenza virus strains. Wells were washed with PBS and then blocked for 1h at 37°C with 200 μl/well of RPMI containing 10% heat-inactivated FBS. Single marrow cells were extracted from both femurs from each mouse as described above, and cells were then washed twice with PBS containing 0.5% BSA and added to the pre-coated plates at 500,000 cells/well, followed by 1:2 serial dilutions down to 6,250 cells/well. The plates were incubated for 4h in a 5% CO_2_-humidified incubator at 37°C. Thereafter, the wells were washed six times with 0.1% Tween-20 PBS (PBST), and phosphatase-labeled goat-anti-mouse IgG detection antibody was added at 100 μl/well. Plates were then incubated overnight at 4°C, washed six times with PBST, before IgG-specific spots were developed using the Alkaline Phosphatase Substrate Kit III. Spot numbers were enumerated using an ImmunoSpot plate reader (Cellular Technology Limited; Cleveland, OH, USA).

### Flow cytometry and cell sorting

Leukocytes were harvested from spleen as well as inguinal and mediastinal lymph nodes by passage through a 40-μm mesh and incubated for 10 min with 1 μg/mL of anti-CD16/32 Fc-block in FACS buffer (PBS supplemented with 1 mM EDTA, 2% heat-inactivated FBS, and 0.02% sodium azide). Cells were further labeled using fluorochrome-coupled antibodies directed against the respective antigens: CD45R, GL-7, CD95, CD138, IgG, IgG1, IgG2c, CXCR4, CD83, CD4, PD-1, CXCR5, CD77, Gr-1, F4/80, CD103, and CD11b. Direct detection of antigen-binding B cells was performed using NP-PE and OVA-FITC labeling of singe cell suspensions. For phospho-Akt staining, B cells were stimulated with anti-BCR antibodies for precisely the same time to avoid variations induced by intracellular phosphatases. Cells were surface-stained at 4°C for 30 minutes, followed by fixation and permeabilization using the Fixation/Permeabilization buffer kit according to manufacturer’s instructions (BD Biosciences, NJ, USA). This was followed by staining with anti-phospho-Akt antibody diluted in Perm/Wash buffer for 1h. All flow cytometry data were generated using a FACS Canto Flow Cytometer (BD Biosciences, NJ, USA). For sorting GC B cells, B cells were enriched using a B Cell Isolation Kit (Miltenyi Biotec, Bergisch Gladbach, Germany) from spleens of immunized mice, and subsequently, GC B cells (CD45R^+^, GL7^+^, CD95^+^) and naïve B cells (CD45R^+^, GL7^-^, CD38^-^) were sorted either using a FACS Aria II (Becton Dickinson) or SH-800 cell sorter (Sony).

### Confocal microscopy

Confocal microscopy was performed to visualize germinal centers as well as the results of proximity ligation assays. Regarding the former, spleens from immunized mice were fixed in 4% paraformaldehyde in PBS for 1h at 4°C, followed by a five-time wash in PBS (10 minutes each). Thereafter, spleens were moved to 30% sucrose solution in PBS (overnight) and were frozen in TAK tissue-mounting media the following day. For immunostaining, 30-μM sections were cut, air dried for 1h, prior to rehydration in PBS with 1% BSA for 10 min. Slides were then washed three times in PBST and blocked for 1h at room temperature with 2% BSA. Staining with primary antibodies directed against CD3, GL7, and IgD was performed in PBS with 2% mouse serum, 0.1% BSA, and 0.1% NaN_3_ at 4°C overnight. Subsequently, slides were mounted with anti-fade mounting medium, and images were taken using an Olympus Fluoview (FV) 1000 confocal microscope.

### Proximity ligation assay

FACS-sorted GC B cells were plated on PTFE-slides for 30 min at 37°C and were fixed for 15 min with 2% paraformaldehyde. For intracellular PLA, cells were permeabilized after fixation with 0.5% saponin in PBS for 30 min. PLA was carried out with Duolink® in situ detection reagents following the manufacturer’s instructions (Millipore Sigma; MA, USA). Briefly, fixed cells were blocked using a solution that contained 25 μg/ml sonicated salmon sperm DNA and 250 µg/ml BSA in PBS. After blocking, cells were incubated with appropriate PLA probes directed against either CD19, BCR, CD81, or PI3K, and cells were washed with PBS thrice. Thereafter, cells were further incubated with secondary PLA probes binding to the corresponding primary antibodies. To study the interaction between Gb3 and CD19, anti-mouse CD19 and anti-Gb3 antibodies were directly tagged to plus and minus oligo probes, using the PLA probe maker kit following the manufacturer’s guidelines (Millipore Sigma MA, USA), and fixed cells were directly labeled with tagged antibodies overnight. Subsequently, all probe-labeled samples were subjected to PLA signal amplification following the manufacturer’s instructions. Finally, cells were mounted on slides with DAPI-containing mounting reagent. Images were taken using an Olympus Fluoview (FV) 1000 confocal microscope, and data were analyzed using ImageJ software (NIH).

### Isothermal titration calorimetry (ITC) assay

ITC experiments were performed using a MicroCal ITC microcalorimeter at 25°C and a 150 mM NaCl buffer at pH 7.4. Recombinant CD19 was dialyzed into PBS containing 20 mM HEPES, and the lyophilized Gb3 analog was suspended in dialysate to achieve buffer match. All experiments were performed with the CD19 ectodomain in the microcalorimeter cell at 300 µM, and the Gb3 analog in the syringe at 100 mM. The titrations consisted of a total of 35 injections (5 µl each), spaced 80 seconds apart. Protein concentrations were determined based on the UV absorbance at 280 nm, and the molar extinction coefficient of the CD19 ectodomain was 6,805 M^−1^cm^−1^. Data were analyzed based on MicroCal ITC Origin 7 software (Malvern Panalytical Ltd, Worcestershire, UK) using a one-site binding model.

### Western blotting

FACS-sorted GC B cells were lysed using RIPA buffer containing 1x protease and phosphatase inhibitors, and the protein concentrations of whole cell lysates were quantified using the BCA protein assay kit. The samples were boiled in SDS loading Laemmli buffer (BioRad) supplemented with β-mercaptoethanol and subsequently resolved using a 12% polyacrylamide gel, prior to transfer onto a nitrocellulose membrane. Next, the membrane was blocked with 5% BSA for 1h and then probed for the indicated proteins using specific primary antibodies overnight. All primary antibodies were used at 1:500, which were further probed with HRP-labeled donkey anti-rabbit antibody (Jackson Immuno-Research, West Grove, PA, USA). HRP-labeled antibody/protein complexes were detected by incubating the membrane with enhanced chemiluminescent substrate and exposing the blot to X-ray films in the dark.

### Protein lipid overlay assay

Protein lipid overlay assays were performed as previously described (*51*). Various glycolipids were dissolved in a mixture of methanol and chloroform (1:1), spotted onto a PVDF membrane and air-dried. The membrane was further blocked with 4% fatty acid-free BSA in PBST buffer (PBS containing 0.1% Tween 20, pH 8.0) for 1h at room temperature. Thereafter, the membrane was incubated with histidine-tagged recombinant CD19 (5 μg/ml) at room temperature for 2h. Subsequently, the membrane was washed 5 times (5 min each) with PBST buffer and incubated for 1h at room temperature with a 1:4,000 dilution of rabbit anti-histidine antibody. Next, the membrane was washed 5 times as described above and then incubated with a 1:10,000 dilution of HRP-labeled donkey anti-rabbit antibody. Finally, the membrane was washed in PBST buffer, and deposited lipids were detected by enhanced chemiluminescence on X-ray films.

### Lipid loading onto GC B cells

GC B cells were isolated from Gb3-deficient *A4galt*-KO mice and purified as described above. For lipid incorporation, a complex of Gb3 or its analogs with defatted BSA was prepared in serum-free RPMI supplemented with 15 mM HEPES (pH 7.2) at a molar ratio of 1:1. Cells were seeded in V-shaped 96-well plates and incubated with the lipid/BSA solution for 2h at 37°C. Subsequently, cells were washed once with complete RPMI/FCS and once with PBS, prior to collection by centrifugation and resuspension in complete RPMI/FCS for further stimulation assays. Gb3 incorporation into B cells was confirmed by flow cytometry using anti-Gb3 antibody staining.

### BCR sequencing

Splenic naive and GC B cells were enriched using a B cell isolation kit followed by FACS-sorting as described above. Rep-SHM-Seq was performed as described previously using our established bioinformatics pipeline (*18*). We used bait primers close to the coding ends of JHs to capture full-length V(D)J sequences for SHM analysis and used Illumina MiSeq 2 × 300-bp paired-end sequencing (*18, 52*). As described before, mixed JH primers were designed based on a highly degenerate region to minimize the amplification bias caused by different primers. 400 ng genomic DNA from FACS-sorted naive or GC B cells was used as starting material to keep the level of PCR amplification comparable among different samples. In addition, to minimize the potential cross-contamination due to FACS sorting, we further filtered reads by assigning mutated sequences to GC B cell samples and non-mutated sequences to naive B cell samples. Libraries for sequencing were prepared as described(*17*). Briefly, genomic DNA was sonicated and subjected to linear amplification-mediated PCR using biotinylated bait primers. Linear amplification-mediated PCR products were purified using Dynabeads MyOne streptavidin C1 magnetic beads (Thermo Fisher Scientific, MA, USA) and ligated to bridge adaptors. Adaptor-ligated products were amplified by nested PCR with indexed J_H_1/J_H_3 primers as well as primers annealed to the adaptor. PCR products were further tagged with Illumina sequencing adaptor sequences and size-selected via gel extraction. Libraries were sequenced by paired-end 300-bp sequencing on MiSeq, an Illumina sequencing platform (San Diego, CA, USA).

#### a. V segment usage analysis

For each sample, the ratio of each functional V segment was calculated as % usage among total productive and non-productive V(D)J junctions. For GC vs naive repertoire comparison, V segment usages from multiple samples were plotted in the same chart above and below the x-axis along indicated chromosomal coordinates. A complete list of functional V_H_ segments in the order plotted along the x-axis is shown in Table S1. Significantly enriched V segments in GCs were identified by paired Student’s t-test with multiple test correction by false discovery rate (FDR) and GC-enriched V_H_ segments with FDR value ≤ 15% were considered significant.

#### b. CDR3 clonotype analysis

To identify the enrichment of antigen-specific GC clonotypes across different samples, we pooled the GC productive reads from all the mice of different genotypes together to carry out clonal clustering as described (*18*). In our study, a clonotype is considered as enriched in one sample if it makes up more than 0.3% of all productive junctions. The consensus CDR3 DNA sequence plot for each clonotype was generated by WebLogo (*53*). Clonotype diversity analysis was done by rarefaction using the R package iNEXT. The number of clonotypes was calculated for each sample at the number of reads from 100 to 5000. Student’s t-tests were used to test the difference in diversity across different genotypes at 1000, 3000, and 5000 reads.

#### c. SHM analysis

Mutation frequency in the heavy chain variable segment of CDR3 was calculated by enumerating the number of nucleotide mismatches compared to the background intrinsic non-productive allele for the selected mutation as described (*18*). For statistical analysis, we used Kruskal-Wallis H test with Dunn’s multiple comparison test to compare SHM rate of productive sequences in a clonotype across different genotypes.

### Microneutralization and hemagglutination inhibition assays

Serum samples were pretreated with receptor-destroying enzyme overnight at 37°C, before being heated at 56°C for 30 min and diluted with minimum essential medium to a final serum dilution of 1:10. Serum antibody titers were measured by hemagglutination inhibition (HI) and microneutralization (NA; neutralizing antibody) assays based on standard procedures. For HI assays, we used U-bottom 96-well microtiter plates. Two-fold serial dilutions of sera (25 μl) starting at 1:10 were diluted with PBS and mixed with an equal volume of 4 hemagglutinating units of either H1N1 or H3N2 virus. The mixture of diluted serum and virus was incubated for 30 min at room temperature. Fifty microliters of 0.5% turkey red blood cells were added to the antigen/serum mixture and incubated for 45 min at room temperature. HI antibody titers were expressed as the reciprocal of the highest serum dilution that could prevent hemagglutination. For the NA assay, 50 μl of serial 2-fold dilutions of the sera were prepared and subsequently incubated with equal volumes of 2 x 10^3^ TCID 50/ml of either H1N1 (PR8) or H3N2 (HK68, X-31) for 1 hour at 35°C. Thereafter, 1.5 x 10^4^ MDCK cells in 96-well plates were incubated with serum/virus mixtures at 35°C for 72 hours. 50 μl of culture supernatant from each well was incubated with 50 μl of 1% turkey red blood cell in 96-well V bottom plate for 1 hour at room temperature. NA titers were defined as the highest dilution factor which can inhibit the infection of MDCK cells with virus.

### Quantification of viral load in the lung

At various time points after infection, lungs were removed and stored at −80°C until virus quantification was performed. Lungs were homogenized individually in 1 ml RPMI. Virus was titrated in duplicate from lung homogenates by standard plaque assay on MDCK cells, and titers were reported as plaque forming units (PFU) per ml RPMI. Briefly, one day before infection, MDCK cells were seeded in six-well plates. The next day, the cells were washed once with 2 ml DMEM and incubated with virus diluted in DMEM containing 2% bovine serum albumin (BSA), 200 U/ml penicillin and 200 ug/ml streptomycin for 1h at 35°C. After incubation, the virus inoculum was removed, and the cells overlaid with DMEM containing 1.6% agarose solution and 2 μg/ml L-(tosylamido-2-phenyl) ethyl chloromethyl ketone (TPCK)-treated trypsin (Sigma). The plates were incubated at 35°C for 48 hours. Subsequently, the plaques were visualized by staining with crystal violet.

### Human tonsil organoids

Dissociated cells from human tonsils were procured from BioIVT (NY, USA) and stimulated with antigens as described previously (*42*). For culture of cryopreserved cells, frozen cells were washed twice in complete medium (RPMI with GlutaMAX, 10% FBS, 1 × nonessential amino acids, 1 × sodium pyruvate, 1 × penicillin/streptomycin, 1 × Normocin from InvivoGen), and 2 × 10^7^ cells per ml were suspended in complete RPMI. 100 μl of cell suspension was then plated per well onto permeable (0.4-μm pore size) membranes (24-well size polycarbonate membranes in standard 12-well plate), with the lower chamber consisting of complete medium (1 ml for 12-well plates, 200 μl for 96-well plates) supplemented with 1 μg/ml of recombinant human B cell-activating factor (BAFF, BioLegend). 1 µg/ml rHA from H1N1 (PR8) was then added directly to the cultured tonsil dissociated cells. For adjuvant testing, either alum (0.01%) or Gb3 (10 µg/ml) were added directly into the culture immediately after antigen addition. Cultures were incubated at 37°C and 5% CO_2_ humidity up to 21 days.

### Statistical analysis

For statistical analysis, samples were randomly selected. Data distribution was checked using Shapiro-Wilk test for normality, and thereafter, analysis for multiple comparison was carried out using Kruskal-Wallis H test with Dunn’s multiple comparison test. To compare two groups, we used either Mann-Whitney U test or Student’s t-test. Student’s t-tests were used to analyze the difference in B cell diversity across different mice, while paired Student’s t-test with multiple test correction by false discovery rate (FDR) was used to determine the enrichment of V_H_ segments in GC B cells compared to naïve B cells. Statistical analyses were performed using Prism (version 9, GraphPad Software) and R version 3.6.3.

## Supplementary figure legends

**Fig. S1.**
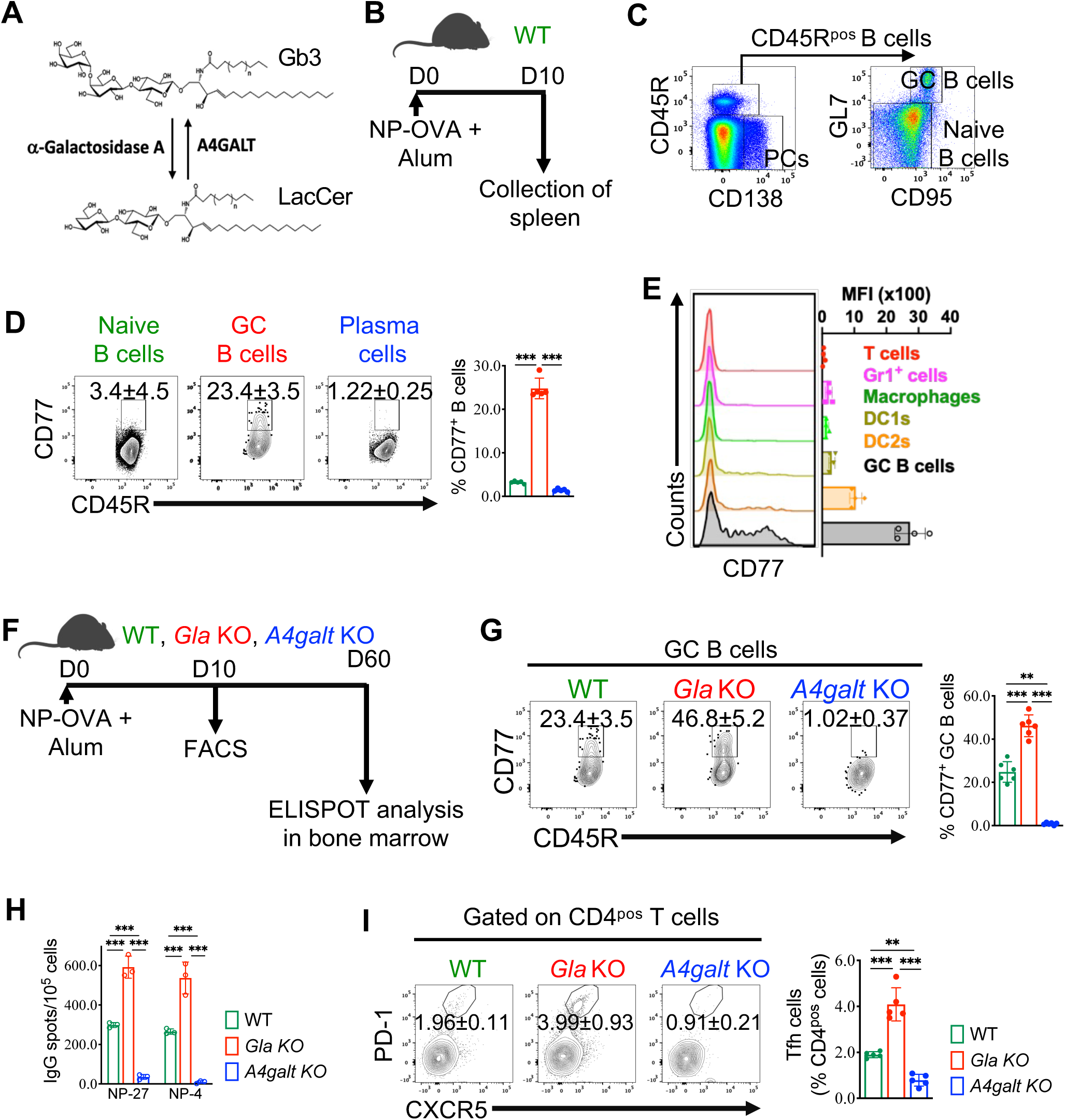
Expression pattern of Gb3 in cell lineages, modulation in enzyme-deficient mouse models, and differentiation of Tfh and plasma cells. (A) Enzymatic pathways of the synthesis and degradation of Gb3. (B) Experimental set-up. WT mice were immunized with NP-OVA adsorbed on alum, and spleens were harvested 10 days later. (C) Gating strategy to identify splenic B cell subsets by flow cytometry. (D) Representative FACS plots and percentages of Gb3-positive B cells on day 10 post NP-OVA immunization. Each dot represents one mouse (5 mice per group), and the experiment was repeated three times. (E) Representative histogram overlays depicting Gb3 expression on splenic myeloid and lymphocyte subsets on day 10 post immunization quantified by flow cytometry. DC1 = XCR-1^+^ classical DCs, DC2 = CD11b^+^ classical DCs, Gr1^+^ cells = neutrophils/monocytes. Each dot represents a single mouse (n=4), and the experiment was repeated twice. (F) Schematic representation of the experimental set-up. WT, *Gla*-KO, or *A4galt*-KO mice were immunized with NP-OVA adsorbed on alum. (G) Representative FACS plots and percentages of Gb3-positive GC B cells on day 10 post NP-OVA immunization. Each dot represents one mouse (6 mice per group), and the experiment was repeated twice. (H) Quantification of plasma cells secreting NP-specific total IgG and NP4-specific high-affinity IgG by ELISPOT. Each dot represents one mouse (3 mice per group), and the experiment was repeated at least three times. (I) Representative FACS plots and percentages of Tfh cells in the spleen on day 10 post NP-OVA immunization were quantified by flow cytometry. Each dot represents one mouse (5 mice per group), and the experiment was repeated at least three times. The *p*-values in all the data sets were calculated by Kruskal-Wallis H test with Dunn’s multiple comparison test.

**Fig. S2.**
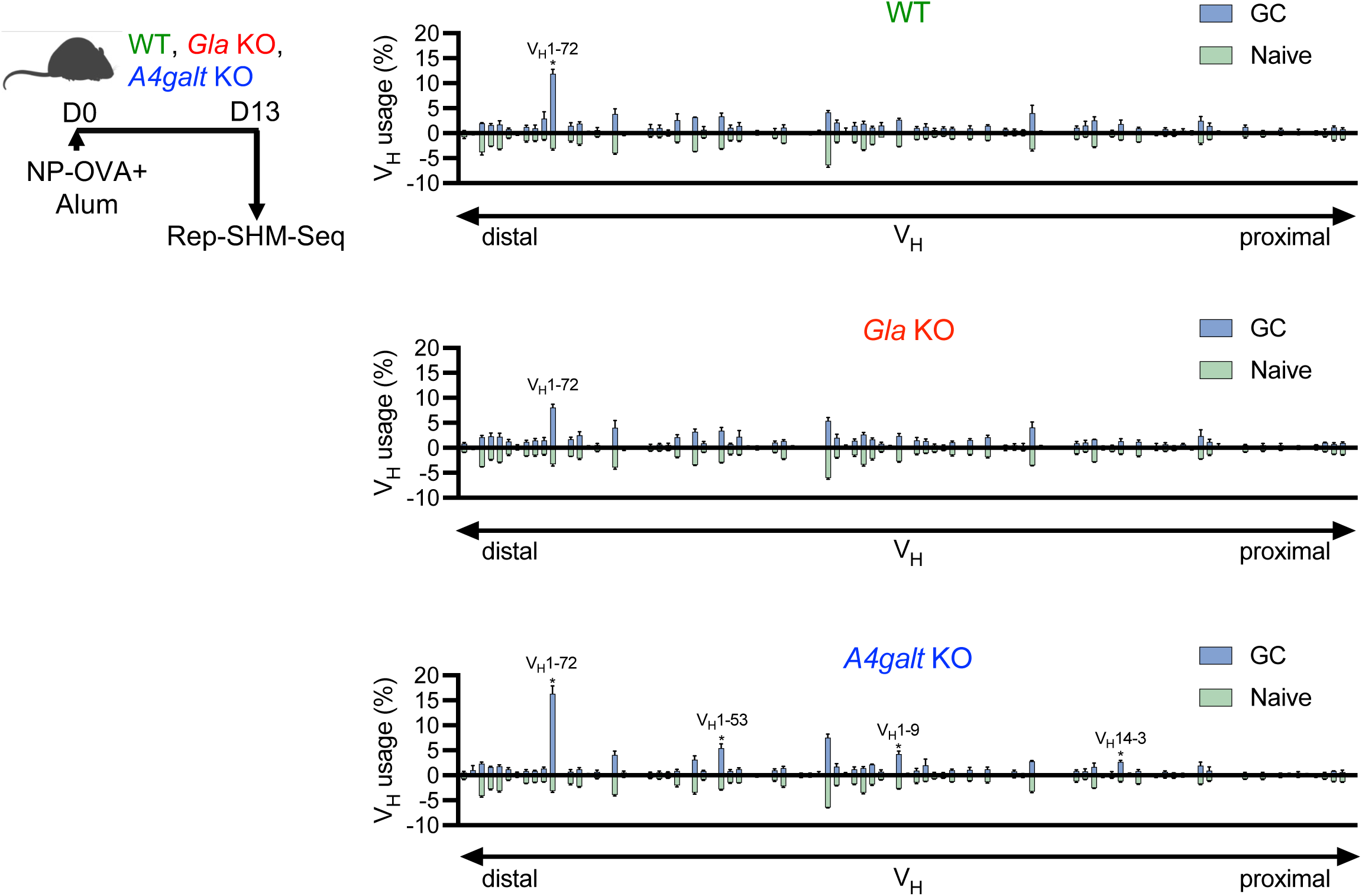
Gb3 abundance does not affect V_H_ usage in germinal centers. Shown is the V_H_ repertoire of productive V(D)J junctions in splenic GC B cells versus naive B cells in WT, *Gla*-KO, or *A4galt*-KO mice immunized with NP-OVA. The data are plotted as mean ± SD (N=3 for WT and *Gla*-KO, and N=4 for *A4galt*-KO). V_H_ segments are each ordered linearly based on their chromosome coordinates with respect to relative proximity to the D segments. Paired Student’s t-test was performed to compare BCR usage between GC and naïve B cells, and GC-enriched V_H_ segments with FDR value ≤ 15% were marked with *, along with a usage cut-off over 1%.

**Fig. S3.**
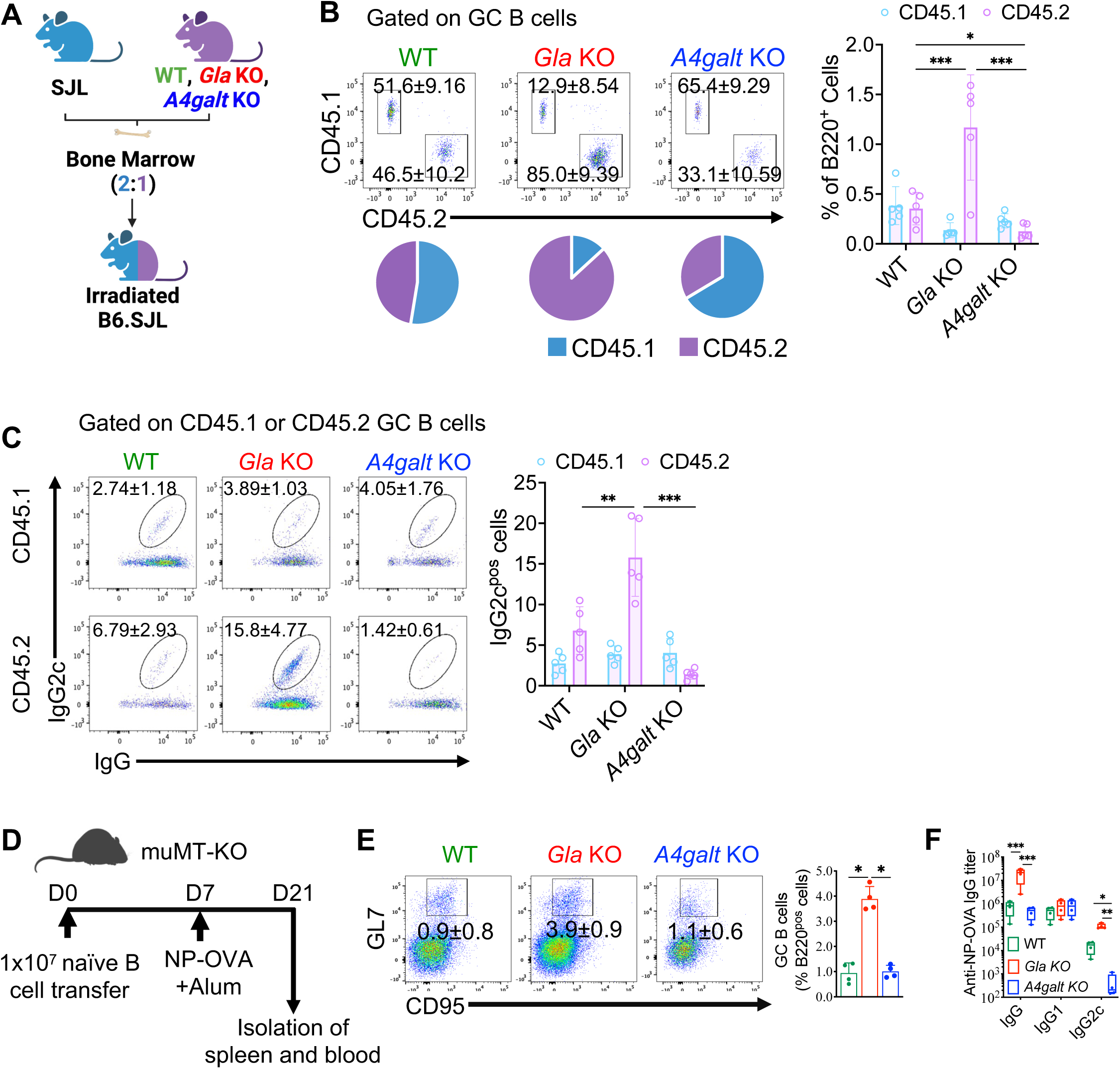
Gb3 abundance on GC B cells provides competitive advantage during GC reaction in a B cell-intrinsic manner. (A) Experimental scheme depicting generation of congenic BM chimeric mice. WT.SJL recipient mice were lethally irradiated and intravenously transferred with WT CD45.1 (SJL) and CD45.2 (C57BL/6, *Gla*-KO, or *A4galt*-KO) BM cells at a ratio of 2:1. (B) Representative FACS plots and relative percentages of CD45.1^+^ and CD45.2^+^ GC B cells in the spleen on day 10 post NP-OVA immunization. Each dot represents one mouse (5 mice per group), and the experiment was repeated twice. (C) Representative FACS plots and relative IgG2c-positive CD45.1^+^ and CD45.2^+^ GC B cells in the spleen on day 10 post NP-OVA immunization. Each dot represents one mouse (5 mice per group), and the experiment was repeated twice. (D) Experimental set-up showing adoptive B cell transfer and NP-OVA immunization strategy. muMT-KO mice were injected with 1 × 10^7^ B cells intravenously and were immunized on day 7 with NP-OVA adsorbed on alum. (E) Representative FACS plots and percentages of GC B cells in the spleen. Experiment was repeated twice (n=4 mice per group). (F) Serological analysis of immunoglobulin concentrations of indicated isotypes as detected by end-point titer measurements using ELISA on day 21 after immunization. Experiment was repeated twice with 4-5 mice per group. The *p*-values were calculated by Kruskal-Wallis H test with Dunn’s multiple comparison test.

**Fig. S4.**
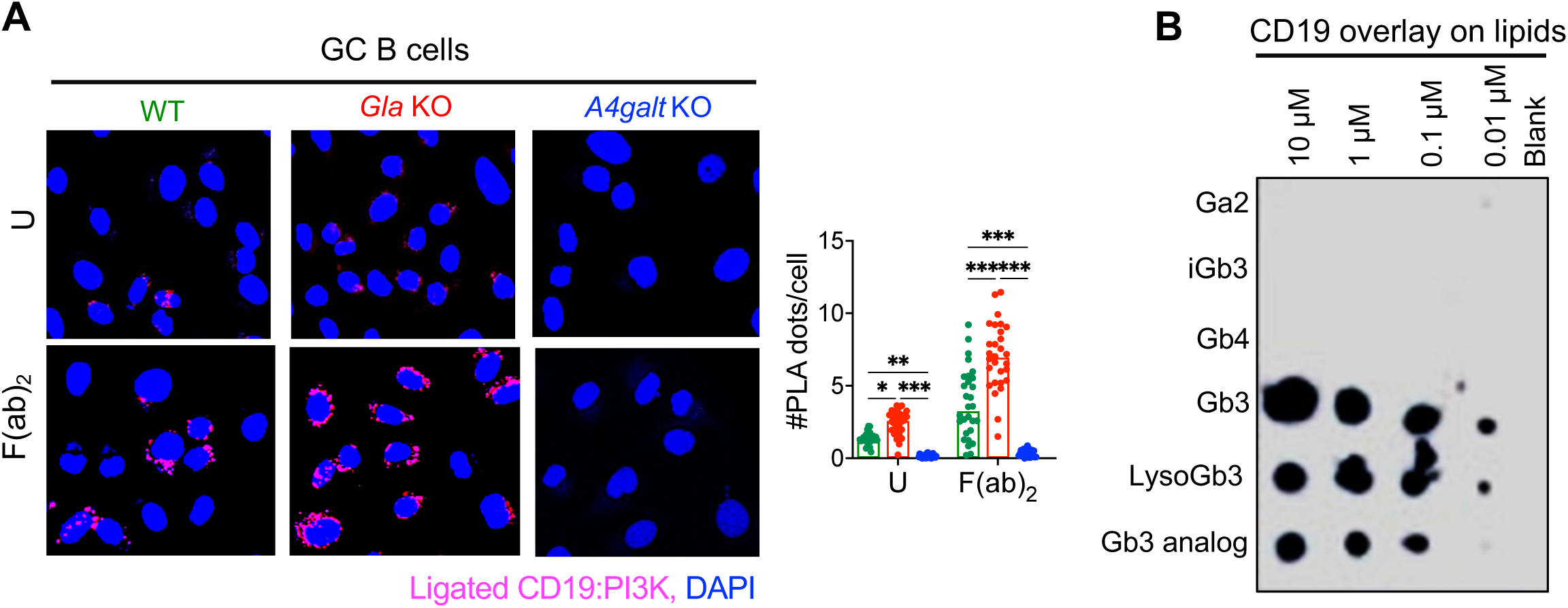
Gb3 binds to CD19 and recruits PI3K. (A) Proximity ligation assay (PLA) performed on GC B cells to quantify the recruitment of PI3K to CD19 (blue = DAPI, red = CD19:PI3K PLA signal). FACS-sorted GC B cells were stimulated with anti-BCR antibodies (F(ab)_2_) and were probed with specific antibodies. Experiment was repeated three times, and PLA signal on more than 30 cells in different fields was calculated for statistical analysis. U = unstimulated. (B) Protein lipid overlay assay to assess the CD19 binding capability of different lipids. Lipids were deposited on a PVDF membrane and their binding to CD19 (containing histidine) was tested by using anti-His-HRP secondary antibody using chemiluminescence. The *p*-values in graph were calculated by Kruskal-Wallis H test with Dunn’s multiple comparison test.

**Fig. S5.**
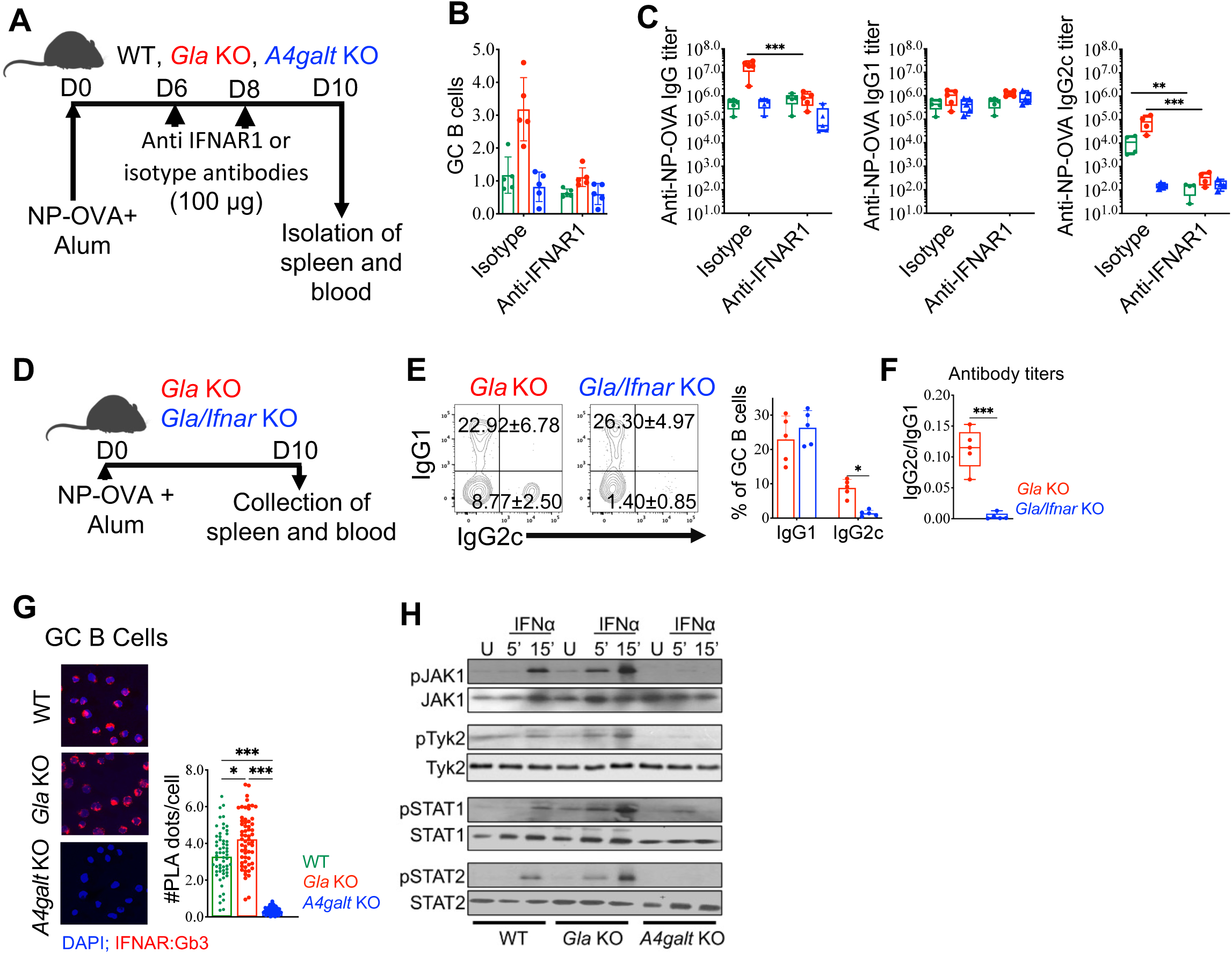
Gb3-mediated IgG2c isotype switch is IFNAR1-dependent. (A) Experimental set-up depicting immunization and anti-IFNAR1 antibody treatment. Mice were immunized with NP-OVA adsorbed on alum, and blocking antibodies were administered on day 6 and 8 after immunization. (B) Frequencies of GC B cells in the spleen on day 10 post immunization. Each dot represents one mouse (4 mice per group), and the experiment was repeated at least three times. (C) Anti-NP-OVA immunoglobulin titers of indicated isotypes in the serum were measured by ELISA. Each dot represents one mouse (4 mice per group), and the experiment was repeated at least three times. (D) Experimental set-up depicting NP-OVA immunization of *Gla*- and *Ifnar*-deficient mice. (E) FACS plots and percentages of IgG1^+^ and IgG2c^+^ GC B cells in the spleen were quantified by flow cytometry. Dots in bar graphs depict individual mice (n=5 mice per group). (F) Ratio of IgG2c to IgG1 titers in the serum. Bar graph shows the ratio of endpoint titers for IgG2c and IgG1, and each point in the graph depicts an individual mouse (n=5 mice per group). (G) PLA performed on GC B cells to probe for proximity between Gb3 and IFNAR1. Images were captured by confocal microscopy (blue = DAPI, red = IFNAR:Gb3 PLA signal). Experiments were repeated at least three times, and PLA signal on more than 30 cells in different fields was calculated for statistical analysis. (H) Immunoblot showing IFNAR1 downstream signaling. IFNα was used to stimulate B cells for 5 or 15 minutes. U = unstimulated. The *p*-values in graph B and G were calculated by Kruskal-Wallis H test with Dunn’s multiple comparison test, while two-sided Mann-Whitney U test was used for analysis in graph C, E, and F.

**Fig. S6.**
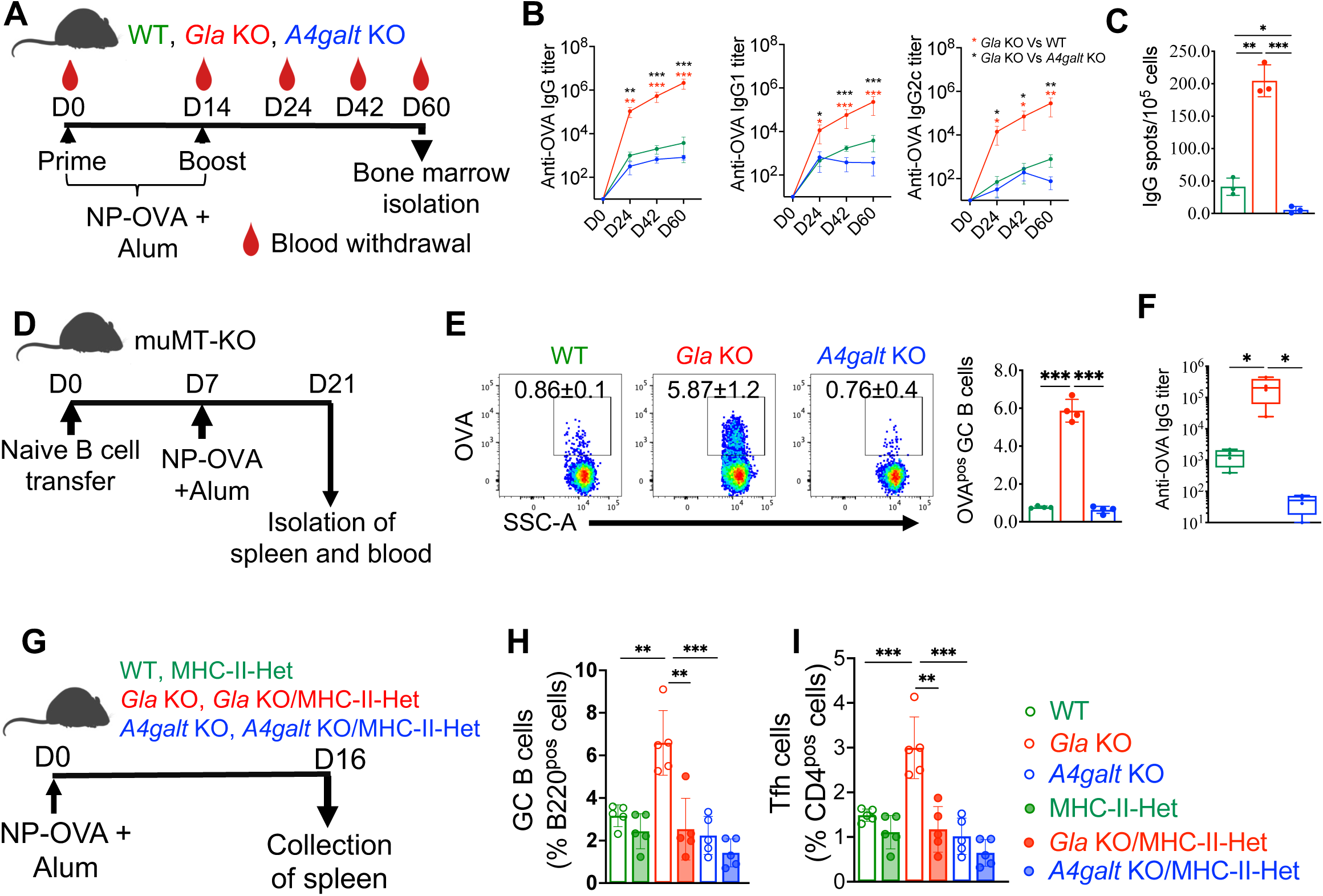
Gb3 facilitates B cell responses to subdominant epitopes. (A) Experimental set-up depicting animal groups, timeline of immunization, and sample collection schedule. (B) Serological analysis of immunoglobulin concentrations of indicated isotypes specific for OVA was performed by ELISA. Graphs represent mean ± SD with 5 mice in each group. The experiment was repeated twice. (C) IgG-producing plasma cells recognizing OVA in the bone marrow on day 60 after immunization using ELISPOT. The experiment was repeated three times (3 mice per group). (D) Experimental set-up showing adoptive B cell transfer and NP-OVA immunization strategy. muMT-KO mice were injected with 1 × 10^7^ B cells intravenously and immunized on day 7 with NP-OVA adsorbed on alum. (E) Representative FACS plots and percentages of OVA-FITC binding by GC B cells in the spleen. Each dot represents one mouse (4 mice per group), and the experiment was repeated at least three times. (F) Concentration of anti-OVA IgG titers in the serum. Experiment was repeated at least three times (n=4-5). (G) Experimental set-up of NP-OVA immunization in MHC-II-haplosufficient mouse strains. (H) Percentage of GC B cells in the spleen on day 10 post NP-OVA immunization. The experiment was repeated three times (5 mice per group). (I) Percentages of Tfh cells in the spleen on day 16 post NP-OVA immunization. Bar graph shows data from individual mice (5 mice per group with experiment repeated at least three times). *p*-values in all the graphs were calculated by Kruskal-Wallis H test with Dunn’s multiple comparison test.

**Fig. S7.**
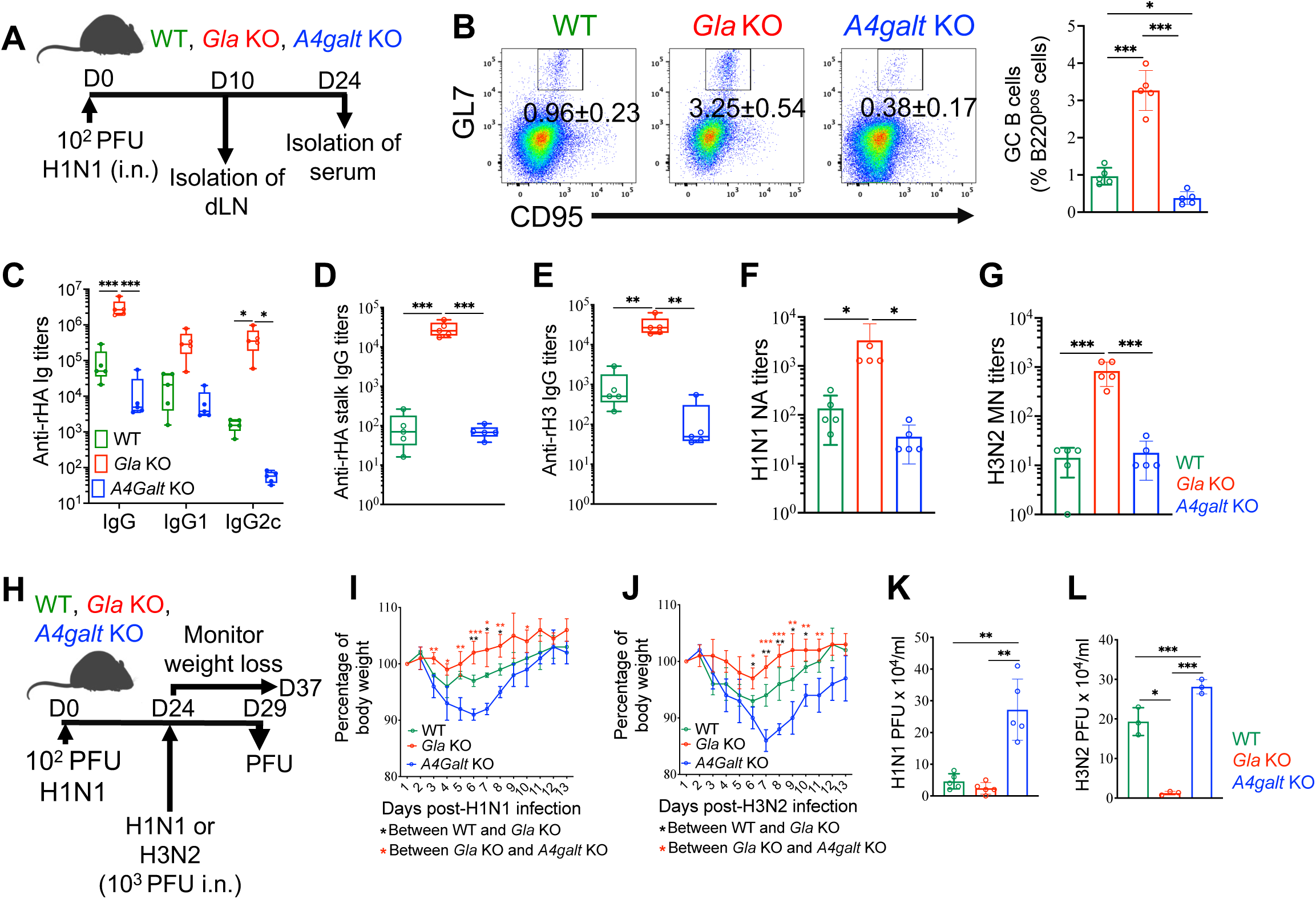
Gb3 drives protection against heterotypic influenza infection. (A) Experimental schedule of H1N1 infection (PR8; 10^2^ PFU) of different mouse strains. PFU = plaque forming units; i.n. = intranasal. (B) Representative FACS plots and percentages of GC B cells in mediastinal draining lymph node. Bar graph depicts individual mice (5 mice per group), and experiment was repeated at least three times. (C) Serological analysis of immunoglobulin concentrations of indicated isotypes as performed by end-point titer measurements using ELISA. Graph depicts individual mice (5 mice per group), and experiment was repeated at least three times. (D) Serum concentration of anti-rHA stalk immunoglobulins was detected by ELISA. Graph depicts individual mice (5 mice per group), and experiment was repeated at least three times. (E) Serum concentrations of immunoglobulins against rH3 (from H3N2) were detected by ELISA. Graph depicts individual mice (5 mice per group), and experiment was repeated at least three times. (F and G) H1N1 and H3N2 neutralizing antibody (NA) titers in serum on day 24 post infection. Graphs depict individual mice (5 mice per group), and experiments were repeated at least three times. (H) Experimental set-up of H1N1 (PR8) or H3N2 (HK68, X-31) influenza infection. H1N1-primed mice were challenged with 10^3^ PFU of either H1N1 or H3N2. PFU = plaque forming units; i.n. = intranasal. (I and J) Weight loss curve following H1N1 or H3N2 infection. Data shown represent three independent experiments with 5 mice in each group. (K and L) H1N1 or H3N2 PFUs in lungs on day 5 post infection. Dots in bar graphs represent individual mice, and the experiments were repeated twice. *p*-values in all graphs were calculated by Kruskal-Wallis H test with Dunn’s multiple comparison test, and in graph I and J, *p*-values of ≤0.05, ≤0.01, and ≤0.001 were marked with *, **, or ***, respectively.

**Fig. S8.**
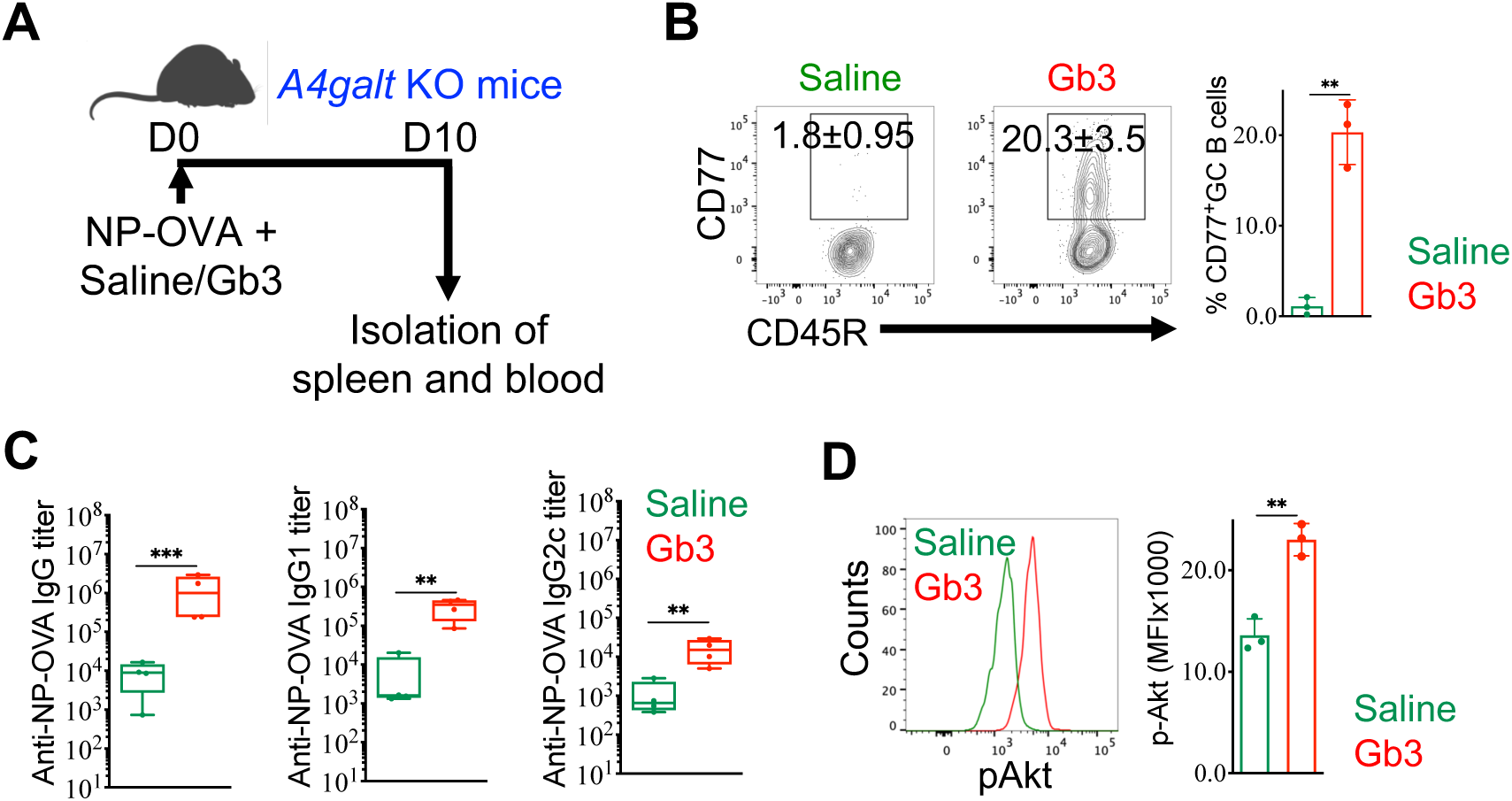
GC B cells acquire exogenous Gb3 upon immunization to deploy its adjuvant functions. (A) Experimental set-up for immunization of *A4galt*-KO mice with NP-OVA in the presence of Gb3. (B) Representative FACS plots and graph showing Gb3-positive GC B cells after immunization of *A4galt*-deficient mice with Gb3. Graph depicts data from 3 mice per group, and the experiment was repeated twice. (C) Serum concentrations of immunoglobulins against NP-OVA were detected by ELISA. Each dot represents one mouse. (D) Representative histogram overlay and bar graph showing the expression of phospho-Akt in GC B cells after BCR stimulation. Each dot represents one mouse, and the experiment was repeated twice. *p*-values for all graphs were calculated by two-sided Mann-Whitney U test.

**Fig. S9.**
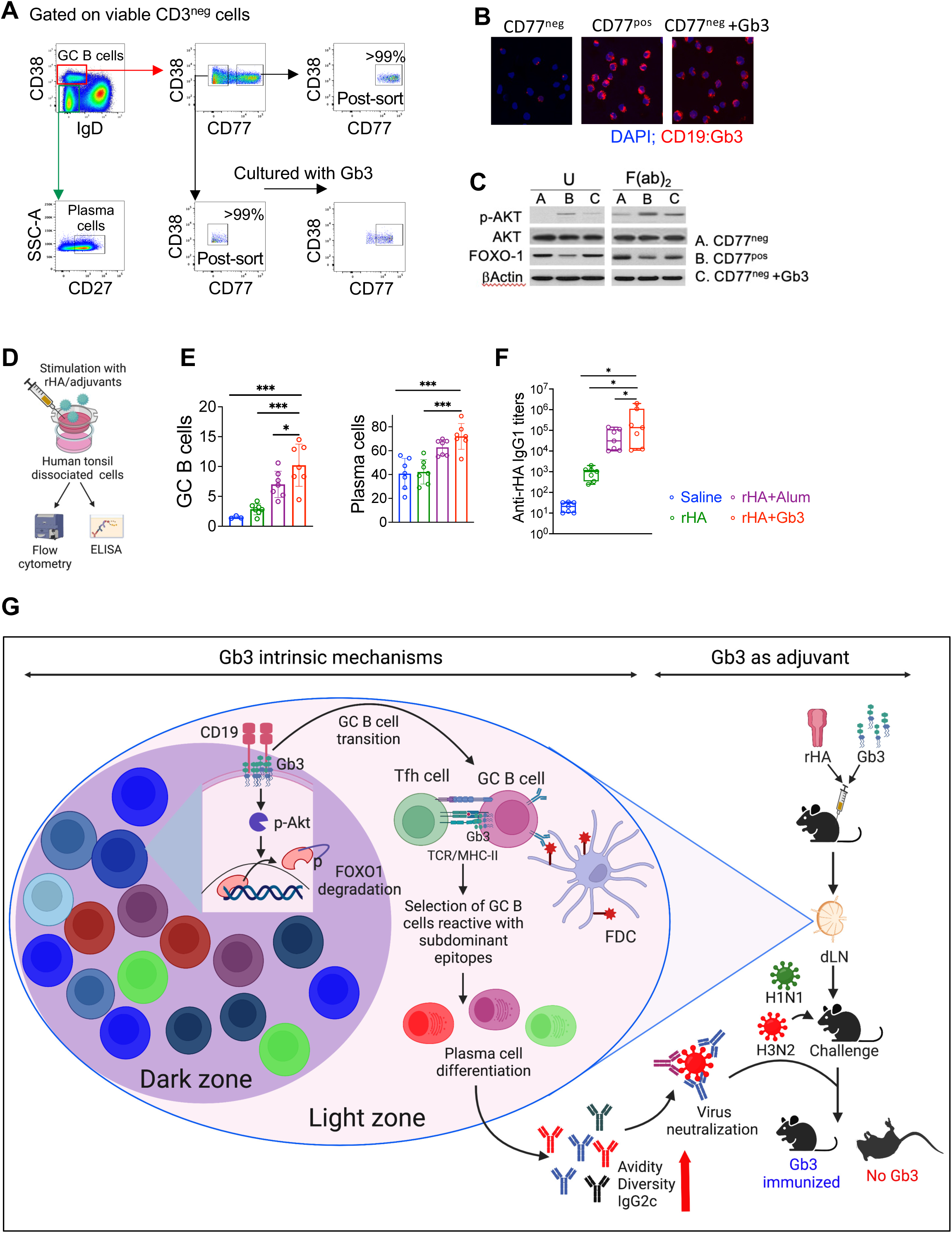
Gb3 interacts with CD19 in human GC B cells to facilitate downstream signaling and serves as adjuvant in human tonsil organoids. (A) Representative FACS plots showing gating strategy to identify B cell subsets among human tonsil cells. GC B cells (CD38^+^); plasma cells (CD27^+^, IgD^-^). CD77-positive and CD77-negative GC B cells were sorted from human tonsils, and CD77-negative GC B cells were subsequently cultured with Gb3. FACS analysis shows Gb3-positivity after culture (bottom panel). CD77 = Gb3. (B) Proximity ligation assay (PLA) performed on GC B cells to probe for vicinity between Gb3 and CD19 (blue = DAPI, red = CD19:Gb3 PLA signal). CD77-negative, CD77-positive, and CD77-negative B cells reconstituted with Gb3 were used for PLA. (C) Immunoblot showing CD19 downstream signaling molecules (Akt) and transcription factors (Foxo1). CD77-negative, CD77-positive, and CD77-negative B cells reconstituted with Gb3 were used for immunoblot analysis. Anti-BCR antibodies (F(ab)_2_) were used to stimulate B cells. U = unstimulated. (D) Workflow for stimulation of human tonsil organoids with antigens and adjuvants. (E) Graphs depicting GC B cell and plasma cell percentages from unstimulated or antigen/adjuvant-stimulated tonsil organoids on day 10 post treatment measured by flow cytometry. Each dot in bar graph represents a separate human tonsil donor (n=7). (F) IgG1 secretion from unstimulated or antigen/adjuvant-stimulated tonsil organoids on day 21 post treatment. Each dot in graph represents a different human tonsil donor (n=7). In all graphs, data were analyzed using Kruskal-Wallis H test with Dunn’s multiple comparison test. (G) Illustration depicts the regulation of the GC B cell response by Gb3 (left panel) and its translation as an adjuvant against influenza infection (right panel). Gb3 binds to CD19 to promote BCR downstream signaling and Foxo1 modulation, driving the cycling of GC B cells to the light zone. Moreover, Gb3 regulates MHC-II antigen presentation to Tfh cells to facilitate selection of antibodies against subdominant epitopes. Use of Gb3 as an adjuvant induces IgG2c class switch and selects antibodies recognizing the stalk region of hemagglutinin. Collectively, these mechanisms provide superior protection against heterologous influenza infection. GC = germinal center; Tfh = T follicular helper cell; TCR = T cell receptor; FDC = follicular dendritic cell; rHA = recombinant hemagglutinin; dLN = draining lymph node; H1N1/H3N2 = influenza virus strains.

---

**Table S1.** V_H_ segment list in the order plotted in Figure S2 (left to right).

| Distal end to D or JH |  |  |  |
| --- | --- | --- | --- |
| 1 | IGHV1-85 | 52 | IGHV1-7 |
| 2 | IGHV1-84 | 53 | IGHV10-3 |
| 3 | IGHV1-82 | 54 | IGHV1-5 |
| 4 | IGHV1-81 | 55 | IGHV1-4 |
| 5 | IGHV1-80 | 56 | IGHV10-1 |
| 6 | IGHV1-78 | 57 | IGHV6-7 |
| 7 | IGHV1-77 | 58 | IGHV6-6 |
| 8 | IGHV1-76 | 59 | IGHV6-4 |
| 9 | IGHV1-75 | 60 | IGHV6-3 |
| 10 | IGHV1-74 | 61 | IGHV12-3 |
| 11 | IGHV1-72 | 62 | IGHV13-2 |
| 12 | IGHV1-71 | 63 | IGHV3-8 |
| 13 | IGHV8-12 | 64 | IGHV9-4 |
| 14 | IGHV1-69 | 65 | IGHV3-6 |
| 15 | IGHV1-67 | 66 | IGHV3-5 |
| 16 | IGHV1-66 | 67 | IGHV3-4 |
| 17 | IGHV8-11 | 68 | IGHV7-4 |
| 18 | IGHV1-64 | 69 | IGHV3-3 |
| 19 | IGHV1-63 | 70 | IGHV14-4 |
| 20 | IGHV1-62-2 | 71 | IGHV7-3 |
| 21 | IGHV1-62-1 | 72 | IGHV9-3 |
| 22 | IGHV1-61 | 73 | IGHV9-2 |
| 23 | IGHV1-59 | 74 | IGHV9-1 |
| 24 | IGHV1-58 | 75 | IGHV14-3 |
| 25 | IGHV8-8 | 76 | IGHV11-2 |
| 26 | IGHV1-56 | 77 | IGHV14-2 |
| 27 | IGHV1-55 | 78 | IGHV11-1 |
| 28 | IGHV1-54 | 79 | IGHV3-1 |
| 29 | IGHV8-6 | 80 | IGHV4-1 |
| 30 | IGHV1-53 | 81 | IGHV14-1 |
| 31 | IGHV1-52 | 82 | IGHV7-1 |
| 32 | IGHV1-50 | 83 | IGHV2-9 |
| 33 | IGHV1-49 | 84 | IGHV5-17 |
| 34 | IGHV1-47 | 85 | IGHV5-16 |
| 35 | IGHV1-43 | 86 | IGHV5-15 |
| 36 | IGHV1-42 | 87 | IGHV2-7 |
| 37 | IGHV1-39 | 88 | IGHV2-6-8 |
| 38 | IGHV1-37 | 89 | IGHV2-9-1 |
| 39 | IGHV1-36 | 90 | IGHV5-12-4 |
| 40 | IGHV1-34 | 91 | IGHV5-9-1 |
| 41 | IGHV1-31 | 92 | IGHV2-6 |
| 42 | IGHV1-26 | 93 | IGHV5-12 |
| 43 | IGHV1-22 | 94 | IGHV2-5 |
| 44 | IGHV1-20 | 95 | IGHV5-9 |
| 45 | IGHV1-19 | 96 | IGHV2-4 |
| 46 | IGHV1-18 | 97 | IGHV5-6 |
| 47 | IGHV1-15 | 98 | IGHV2-3 |
| 48 | IGHV1-12 | 99 | IGHV5-4 |
| 49 | IGHV1-11 | 100 | IGHV2-2 |
| 50 | IGHV1-9 | 101 | IGHV5-2 |
| 51 | IGHV15-2 | (Proximal end to D or JH) |  |

